# Cellular remodeling and JAK inhibition promote zygotic gene expression in the *Ciona* germline

**DOI:** 10.1101/2021.07.12.452040

**Authors:** Naoyuki Ohta, Lionel Christiaen

**Affiliations:** Sars International Centre for Marine Molecular Biology, University of Bergen, Bergen, Norway; Center for Developmental Genetics, Department of Biology, New York University, New York, NY, USA

## Abstract

During development, remodeling of the cellular transcriptome and proteome underlies cell fate decisions and, in somatic lineages, transcription control is a major determinant of fateful biomolecular transitions. By contrast, early germline fate specification in numerous vertebrate and invertebrate species relies extensively on RNA-level regulation, exerted on asymmetrically inherited maternal supplies, with little-to-no zygotic transcription. However delayed, a maternal-to-zygotic transition is nevertheless poised to complete the deployment of pre-gametic programs in the germline. Here, we focused on early germline specification in the tunicate *Ciona* to study zygotic genome activation. We first demonstrate that a peculiar cellular remodeling event excludes localized postplasmic mRNAs, including *Pem-1*, which encodes the general inhibitor of transcription. Subsequently, zygotic transcription begins in *Pem-1*-negative primordial germ cells (PGCs), as revealed by histochemical detection of elongating RNA Polymerase II (RNAPII), and nascent transcripts from the *Mef2* locus. Using PGC-specific *Mef2* transcription as a read-out, we uncovered a provisional antagonism between JAK and MEK/BMPRI/GSK3 signaling, which controls the onset of zygotic gene expression, following cellular remodeling of PGCs. We propose a 2-step model for the onset of zygotic transcription in the *Ciona* germline, which relies on successive cellular remodeling and JAK inhibition, and discuss the significance of germ plasm dislocation and remodeling in the context of developmental fate specification.

## Introduction

During embryonic development, defined transitions in the composition of the cellular transcriptome and proteome govern successive cell fate decisions(Moris *et al*, 2016). Common features of fateful molecular transitions include (1) multilineage priming, whereby multipotent progenitors co-express determinants of distinct and mutually exclusive cellular identities(Nimmo *et al*, 2015; Razy-Krajka *et al*, 2014; Hu *et al*, 1997), (2) *de novo* gene expression, which adds to primed factors and completes fate-specific cellular programs(Wang *et al*, 2019; Graf & Enver, 2009), and (3) cross-antagonisms, whereby competing cellular programs inhibit each other upon mutually exclusive fate choices(Wang *et al*, 2013). Transcriptional control exerts a dominant influence on these molecular transitions. Transcription regulators are thus widespread determinants of cell fate decisions, especially in somatic lineages(Levine & Tjian, 2003; Davidson & Levine, 2008; Levine & Davidson, 2005).

In mammals, early germ cell fate specification is also controlled by signal-mediated induction and transcriptional regulation(Ohinata *et al*, 2009; Tang *et al*, 2016; Mitsunaga & Shioda, 2018; Jostes & Schorle, 2018). By contrast, in other vertebrate species such as zebrafish and *Xenopus*, and in numerous invertebrate species, including the fly *Drosophila*, the nematode worm *C. elegans* and the ascidians *Halocynthia* and *Ciona*, early germ cell progenitors are transcriptionally silent(Nakamura & Seydoux, 2008; Kumano *et al*, 2011; Shirae-Kurabayashi *et al*, 2011). This transcriptional quiescence contributes to keeping germline progenitor cells from assuming somatic fates in response to inductive signals from surrounding cells in early embryos(Robert *et al*, 2015; Lebedeva *et al*, 2018). In these systems, the germline is set aside through unequal cleavages and asymmetric divisions, which segregates somatic lineages from primordial germ cells (PGCs), where transcription remains initially silent.

Early unequal cleavages are coupled with polarized distribution of maternal components including the germ plasm, which carries global transcription inhibitors known in several invertebrate species, such as Pgc (polar granule component) in *Drosophila*, PIE-1 in *C. elegans*(Seydoux & Dunn, 1997; Batchelder *et al*, 1999; Blackwell, 2004; Seydoux & Braun, 2006; Hanyu-Nakamura *et al*, 2008), and Pem-1 in ascidians(Kumano *et al*, 2011; Shirae-Kurabayashi *et al*, 2011). Remarkably, although Pgc, PIE-1 and Pem-1 are unrelated proteins thought to have evolved independently in their corresponding phylogenetic lineages, they all inhibit transcription by blocking phosphorylation of Serine 2 in YSPTSPS-like heptapeptide repeats of the C-terminal domain of the RNA Polymerase II main subunit (RNAPII-CTD), which is necessary for transcriptional elongation(Lebedeva *et al*, 2018).

Consistent with progressive segregation of transcriptional quiescence from the whole egg and early blastomeres to primordial germ cells, Pgc, Pie-1 and Pem-1 gene products are among the maternal components that constitute the germ plasm and progressively segregate to PGCs(Mello *et al*, 1996; Nakamura *et al*, 1996; Yoshida *et al*, 1996). In ascidians, *Pem-1* belongs to a group of so-called postplasmic RNAs that are maternally deposited, accumulate to the vegetal-posterior end of the fertilized egg, and are inherited by the earliest germline progenitor cells, named B4.1, B5.2, B6.3 and B7.6, through subsequent unequal cleavages (Figure 1D; (Sasakura *et al*, 2000; Prodon *et al*, 2007)). Consistent with the dominant effect of RNAPII inhibition by Pem-1, this lineage remains transcriptionally silent until an unknown stage.

**Figure 1.**
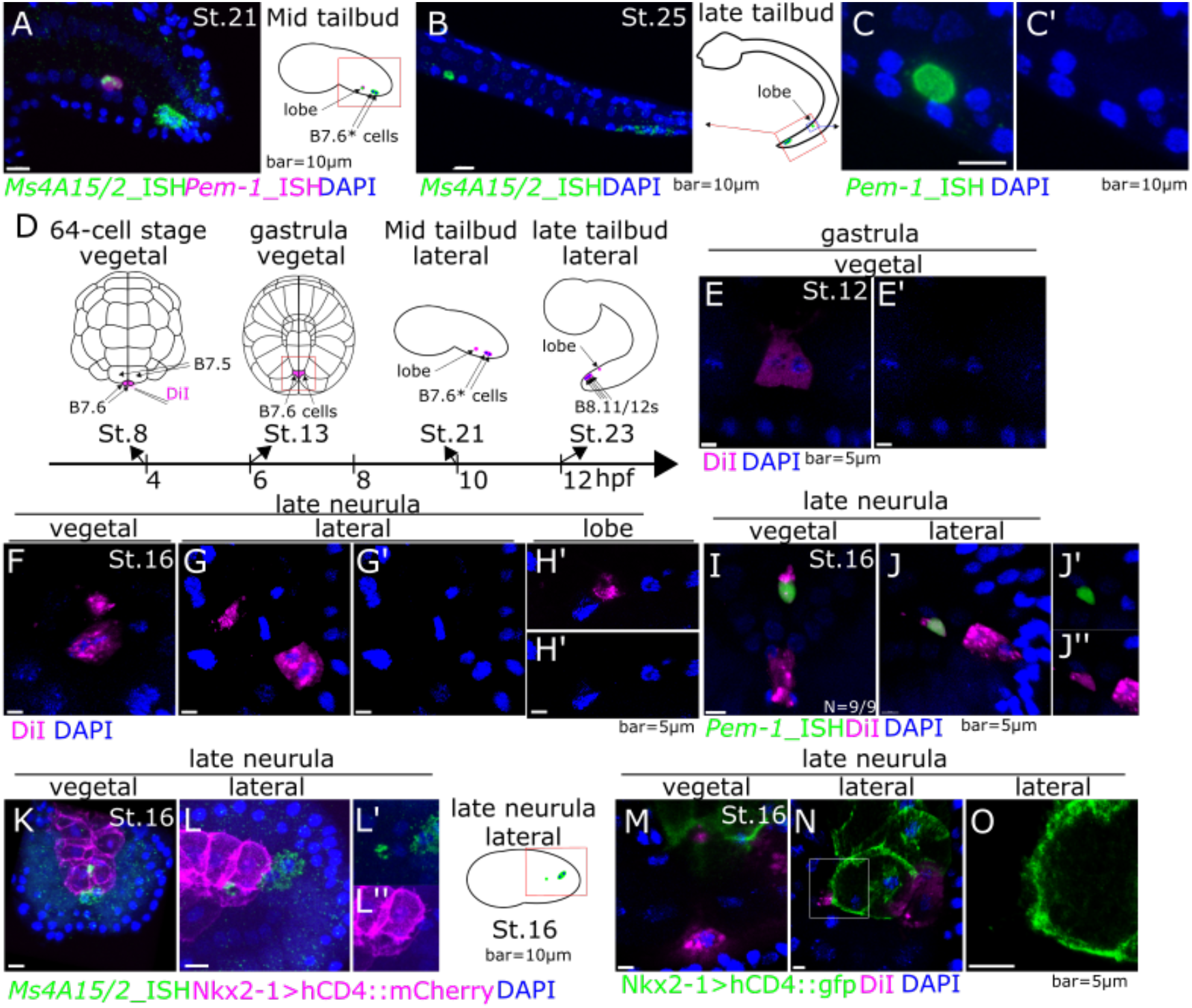
B7.6 cells form lobe including *Pem-1* mRNA without nuclei. (A) Expression of PGC marker genes, *Pem-1* and *MS4a15/2* detected by ISH probes corresponding to exons at the mid tailbud stage. St. 21. (B) Expression of *MS4a15/2* at the late tailbud stage, St. 25. (C, C’) Expression of *Pem-1* in lobe at the late tailbud stage, St. 25. C’ shows the blue channel of C. (D) Schematic diagram of *Ciona* development from 64-cell stage, St. 8 to late tailbud stage, St. 23. Magenta shows DiI label. (E-H’) Cell membranes of B7.6 cells were stained by DiI at the 64-cell stage, and taken image at the gastrula (E, E’) and the neurula stages (F-H’). (E’, G’, H’) Blue channels of E, G and H, respectively. (I-J) B7.6 cells were traced by DiI, and lobe was detected by ISH with a *Pem-1* probe corresponding to exons. (J’) Green and blue channels of J. (J’’) Magenta and blue channels of J. (K-L) Both lobe and B7.6* cells were stained by ISH for *MS4a15/2* gene, and cell membranes of endodermal cells were stained by Nkx2-1>hCD4::mCherry at the late neurula stage, St. 16; ventral (K) and lateral (L) views. (L’) Green and blue channels of L. (L’’) Magenta and blue channels of L. (M-O) Endodermal cell membrane was stained by Nkx2-1>hCD4::gfp in addition to DiI labeling for B7.6 cells. Scale bars=5μm or 10μm.

Remarkably, when B7.6 blastomeres “divide”^***1***^ during gastrulation, *Pem-1* mRNAs are asymmetrically inherited by only one of the “daughter cells”, previously named B8.11, whereas B8.12, its *Pem-1* RNA-negative sibling, constitutes the *bona fide* primordial germ cell (PGC), the progeny of which later populates the somatic gonad in post-metamorphic juveniles(Shirae-Kurabayashi *et al*, 2006). Since *Pem-1* mRNAs are not inherited by PGCs, Pem-1 is likely dispensable for subsequent deployment of the germline-specific program in PGCs.

By contrast with *Pem-1* and several other postplasmic RNAs, mRNAs encoding the Vasa homolog Ddx4, a conserved RNA helicase involved in germ cell development in a broad range of species, are distributed into both *Pem-1+* remnants and the PGCs(Shirae-Kurabayashi *et al*, 2006). Taken together, these observations suggest that maternal determinants of germline fate specification comprise both inhibitors of early somatic specification and primed regulators of the germline program, which segregate upon division of B7.6 blastomeres. We hypothesized that exclusion of *Pem-1* gene products licenses zygotic gene expression in PGCs, thus permitting the activation of *de novo*-expressed factors that complement the germline specification program.

More than 40 maternal RNAs have known postplasmic localization in the zygote and early ascidian embryo(Dehal *et al*, 2002; Yamada *et al*, 2005; Yamada, 2006). By contrast, there is limited-to-no information about zygotically expressed genes in the *Ciona* germline. Contrary to somatic lineages(Imai *et al*, 2006; Ohta & Satou, 2013; Satou & Imai, 2015; Oda-Ishii *et al*, 2016), general transcriptional quiescence has precluded traditional whole genome assays from informing early germline gene regulatory networks (GRNs).

Here, by monitoring the B7.6 lineage in *Ciona* embryos, we first observed that exclusion of *Pem-1* RNAs from the PGCs occurs, not by cell division as previously thought, but through a peculiar cell remodeling event that sheds postplasmic RNA-containing cytoplasm at the beginning of gastrulation. This cellular remodeling is followed by initiation of transcription through the consecutive onsets of RNAPII activity and *Mef2* transcription, at neurula and tailbud stages. Finally, we uncovered a provisional antagonism between JAK and MEK/BMPR/GSK3 signaling that controls the timing of zygotic transcription initiation in the germline. Taken together, these results shed new light on an important transition in early germline development.

## Results

### Cellular remodeling excludes certain maternal postplasmic RNAs from primordial germ cells

In *Ciona* embryos, the B7.6 cells give birth to primordial germ cells (PGCs), which are thought to emerge after one more division, and correspond to the B8.12 lineage, following segregation of a subset of maternal postplasmic RNAs into their B8.11 sister cells(Takamura *et al*, 2002; Shirae-Kurabayashi *et al*, 2006). One of these RNAs, *Pem-1*, produces a nuclear protein that inhibits zygotic transcription in B7.6 cells, thus protecting the PGC lineage by preventing ectopic activation of somatic determinants(Kumano *et al*, 2011; Shirae-Kurabayashi *et al*, 2011). Therefore, we reasoned that zygotic genome activation might follow the exclusion of *Pem-1* RNA from the PGCs. To address this possibility, we first sought to identify candidate zygotically expressed genes, as well as reliable markers of B7.6 lineage cells. We leveraged the extensive *in situ* gene expression database ANISEED(Tassy *et al*, 2010; Dardaillon *et al*, 2020; Brozovic *et al*, 2018). Among genes encoding Postplasmic/PEM RNAs maternally expressed and localized in the B7.6 lineage, we used *Pem-1* (KH.C1.755; (Yamada *et al*, 2005)) as a B8.11 marker, and *Ms4a15/2* (KH.C2.4, aka *Pem-7*; (Nishikata *et al*, 2001; Yamada, 2006)) as a dual B8.11 /B8.12 marker. Whole mount fluorescent in *situ* hybridization (FISH) assays confirmed the expected localization of *Pem-1* and *Ms4a15/2* mRNAs in small B8.11 and large B8.12 cells at the mid tailbud stage (stage 21(Hotta *et al*, 2007); Figure 1A, B). However, to our surprise, we did not find any DAPI-positive nucleus in *Pem-1+* B8.11 “cells” in tailbud embryos (Figure 1C-C’), a pattern visible, but seemingly unnoticed, in a previous publication (Fig.6O in (Shirae-Kurabayashi *et al*, 2006)).

This observation prompted us to reevaluate whether B7.6 cells undergo *bona fide* cell divisions, or cellular remodeling events akin to lobe formation and scission in the primordial germ cells of *C. elegans*(Abdu *et al*, 2016). To this aim, we used cell-specific DiI labeling to monitor B7.6 cell shape changes from the gastrula stage onward (Figure 1D-H’; (Satou *et al*, 2004; Shirae-Kurabayashi *et al*, 2006)). We observed DiI+ cell fragments separate from B7.6 cells (Figure 1F-G’), and lacking DAPI+ nuclei as early as the early neurula stage (Figure 1H-H’). To test whether these B7.6-derived cell fragments correspond to the entities previously recognized as B8.11 cells, we performed fluorescent *in situ* hybridization using the *Pem-1* probe on DiI-labeled neurula stage embryos. This experiment showed colocalization of *Pem-1* RNA with B7.6-derived cell fragments in neurula stage embryos (Figure 1I-J’’). Even though postplasmic RNAs localize the posterior end of the fertilized eggs and early embryos, lobe tended to segregate from the anterior end of B7.6 cells. To clarify this potential conundrum, we inspected embryos collected in time series encompassing lobe formation and scission in embryos. As expected, *Ms4a15/2* mRNAs localized the posterior end of B6.3 cells at 32-cell stage, and the mRNAs were still inherited to the posterior end of B7.6 cells at the beginning of 64-cell stage. During gastrulation however, the FISH signal became visible anteriorly, suggesting a relocalization of postplasmic mRNAs, reflecting changes in B7.6 cell polarity prior to lobe formation (Figure S1A-B). To further probe whether postplasmic RNA exclusion from PGCs occurs via cell division or remodeling, we monitored phosphohistone 3 (pH3) alongside Ms4a15/2 mRNAs to jointly detect mitotic activity and postplasmic RNA localization in B7.6 cells through gastrulation (Figure S1C-D). Clear pH3 signal in dividing B6.3 and some somatic blastomeres confirmed our ability to detect mitotic cells. However, in contrast to a previous report(Shirae-Kurabayashi *et al*, 2006), we observed no pH3 signal in B7.6 nuclei throughout postplasmic RNA repolarization and in the lead up to lobe formation and scission. These observations indicate that B7.6 cells repolarize and undergo a cell remodeling event toward the end of gastrulation, which results in the shedding of cytoplasm containing maternal postplasmic RNA including *Pem-1*, into a cell fragment that we refer to as “lobe”, by analogy with the PGC remodeling process described in *C. elegans*(Mainpal *et al*, 2015; Abdu *et al*, 2016; Maniscalco *et al*, 2020; McIntyre & Nance, 2020). On the other hand, remodeled B7.6 cells, which we propose to call B7.6*, are the *bona fide* primordial germ cells in *Ciona*.

### The endoderm helps PGC remodeling

In various animal species, PGCs associate with endodermal progenitors(Ying *et al*, 2002; Pilato *et al*, 2013). In *C. elegans* for instance, intestinal precursors actively phagocytose the germline lobes(Abdu *et al*, 2016). In ascidians, B7.6 cells abut the posterior-most endodermal progenitors, and the PGCs remain associated with the intestinal anlage, known as endodermal strand, in the larval tail(Nishida & Satoh, 1983; Takamura *et al*, 2002; Kawai *et al*, 2015). We thus explored a possible involvement of the endoderm in PGC remodeling. We combined fluorescent immuno-histochemical (IHC) staining and whole mount *in situ* hybridization to jointly detect the endoderm reporter *Nkx2-1>hCD4::mCherry* (*Nkx2-1* enhancer driving human cluster of differentiation 4 conjugated to monomeric Cherry fluorescent protein)(Ristoratore *et al*, 1999; Gline *et al*, 2015) and the B7.6 lineage marker *Ms4a15/2* in neurula stage embryos. These assays indicated that, while the B7.6 cell bodies remained adjacent to, but outside, the endoderm, the lobe appeared wedged in between endoderm progenitor cells (Figure 1K-L’’). Likewise, joint visualization of endodermal cell membranes, labeled by *Nkx2-1>hCD4::GFP*, and DiI-labeled B7.6 cells revealed the proximity between detached lobes and endodermal progenitors (Figure 1M-O). However, by contrast with *C. elegans*, we did not obtain clear evidence that endoderm cells engulf PGC lobes in *Ciona*. Taken together, these data indicate that B7.6 PGC progenitor cells undergo cellular remodeling events that produce cellular fragments, the lobes, which contain maternally deposited postplasmic mRNAs, and remain in close proximity to endodermal intestine progenitor cells.

### Cellular remodeling precedes the onset of zygotic transcription in PGCs

In ascidians, the maternal postplasmic mRNA *Pem-1* encodes a nuclear protein that inhibits zygotic transcription(Kumano *et al*, 2011; Shirae-Kurabayashi *et al*, 2011). Pem-1 acts by blocking phosphorylation of Serine 2 in heptapeptide repeats at the RNA polymerase II C-terminal domain (RNAPII-CTD)(Kumano *et al*, 2011). We thus reasoned that, by removing *Pem-1* mRNA from B7.6 cells, cellular remodeling may contribute to activating zygotic gene expression in the germline. Indeed, previous immunostaining assays indicated that Pem-1 protein localizes to both nucleus and lobe during B7.6 cell remodeling, but became undetectable in PGCs by the tailbud stage(Shirae-Kurabayashi *et al*, 2011). To test whether the removal of Pem-1 licensed zygotic transcription in remodeled PGCs, we labeled B7.6 cells with DiI, and fixed embryos between the early gastrula and tailbud stages (6 to 12 hours post fertilization (hpf) at 18°C; St. 13, 16, 21 and 23) for immunostaining with an anti-RNAPII-CTD-pSer2 antibody (Figure 2A-D). Consistent with previous reports(Shirae-Kurabayashi *et al*, 2011), we did not detect RNAPII-CTD-Ser2 phosphorylation in B7.6 cells at 6 hpf (St. 13). By contrast, the majority of mid-tailbud stage embryos displayed conspicuous RNAPII-CTD-Ser2 phosphorylation in PGC nuclei (2 nuclei at 10 hpf, and 4 nuclei at 12 hpf after one cell division of B7.6* cells), from 10 hpf (St. 21) onward. These results indicated that transcription elongation by RNA polymerase II is active in the PGCs of mid-tailbud embryos, which follows the exclusion of *Pem-1* mRNAs by B7.6 cell remodeling.

**Figure 2.**
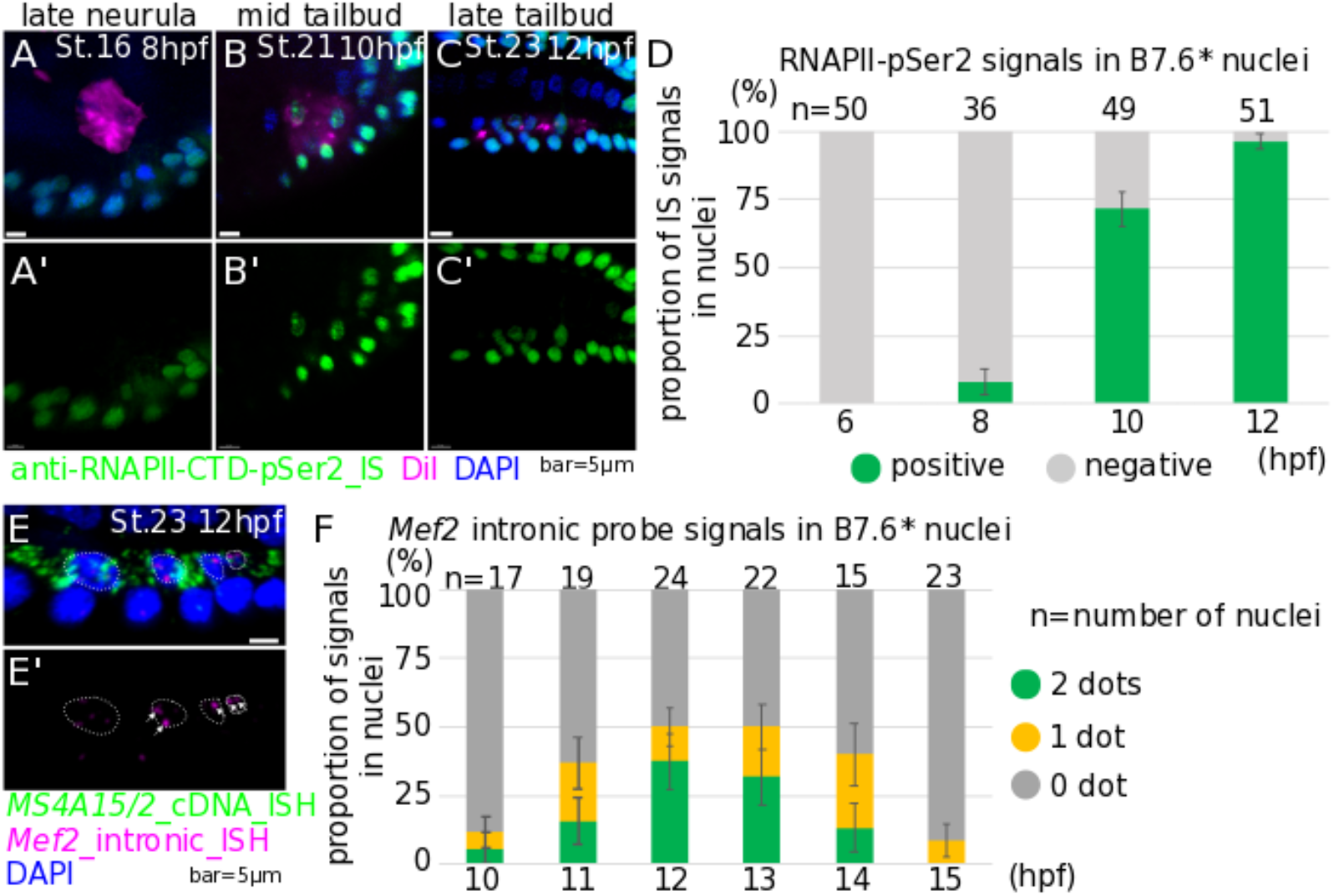
RNA polymerase II Ser2 phosphorylation and *Mef2* nascent expression in B7.6* cells. (A-C) Immunostaining for anti-RNAPII-CTD-pSer2 antibody was done at the late neurula; 8 hpf (A), the mid tailbud; 10 hpf (B) and the late tailbud; 12 hpf (C) stages. (D) Proportion of signal positive nuclei by immunostaining with anti-RNAPII-CTD-pSer2 antibody in B7.6* cells. (E-E’) Nascent expression of *Mef2* gene was detected by *Mef2* intronic probes in 12 hpf embryos at 18 °C. White arrows indicate the dotted signals of nascent *Mef2* expression in nuclei of B7.6* cells. White dotted circles show nuclei of B7.6* cells. (F) *Mef2* intronic probe signals in nuclei in B7.6* cells. Error bars indicate standard error. Scale bars=5μm.

### *Mef2* is zygotically transcribed in the PGCs

Having established that the PGCs of mid-tailbud embryos are transcriptionally active, we sought to identify genes expressed zygotically in B7.6* cells. We mined the gene expression database ANISEED(Tassy *et al*, 2010; Dardaillon *et al*, 2020; Brozovic *et al*, 2018), and identified candidate transcription factor-coding genes possibly upregulated in B7.6* cells after the exclusion of postplasmic RNAs by lobe scission. Here, we focus on a transcription factor coding gene, *Myocyte elongation factor 2* (*Mef2*; KH.S455.6). *Mef2* maternal mRNAs were detected ubiquitously in the whole early embryo and intense ISH signals were observed in the muscle, endoderm and epidermis(Imai, 2004). This abundance of maternal mRNA prevented us from identifying zygotic products with cDNA-derived probes. To specifically monitor zygotic *Mef2* expression, we synthesized *Mef2*-specific intronic probes, which detect nascent transcripts, as shown for *Gata4/5/6, Tbx1/10* and *Ebf* in the cardiopharyngeal lineage(Wang *et al*, 2013; Razy-Krajka *et al*, 2018). To unequivocally identify PGCs, we performed double FISH assays with *Mef2* intronic probe and the B7.6 lineage marker *Ms4a15/2* (Figure 2E). Mid- to late-tailbud embryos raised at 18°C, and collected in a time series between 10 and 15 hpf, St. 21-24, showed conspicuous nuclear dots of nascent transcripts, indicative of zygotic *Mef2* expression in B7.6*, as well as epidermal and muscle cells (Figure 2E-E’). Because transcription was shown to occur in bursts of RNA polymerase activity in other systems(Gregor *et al*, 2014; Tkačik & Gregor, 2021), and our assay provided snapshots of dynamic nuclear states, we reasoned that *Mef2* loci in B7.6* cells might be transcriptionally active and yet show 0, 1 or 2 dots (assuming that even after DNA replication, sister chromatids remain too close to distinguish by standard confocal microscopy). We thus counted *Mef2*+ dots per B7.6* nucleus at successive stages, to obtain a semi-quantitative view of transcriptional activity at the *Mef2* locus in developing PGCs. This analysis indicated that zygotic *Mef2* expression peaked at the late tailbud I stage (st. 23, 12 hpf at 18°C), from an onset around 10 hpf, and was markedly down-regulated by 15 hpf (Figure 2F).

At its peak, we detected 1 or 2 fluorescent dots in 50% of the nuclei, and thus sought to further verify that these correspond to active *Mef2* transcription. We treated embryos with known chemical inhibitors of transcription and assayed zygotic *Mef2* expression with our intronic probe. Actinomycin D, an established transcription inhibitor that binds “melted” DNA at the pre-initiation complex and prevents RNA elongation(Sobell, 1985), was previously used on ascidian embryos(Shirae-Kurabayashi *et al*, 2006; Miyaoku *et al*, 2018), and caused a modest but significant downregulation of *Mef2* expression in B7.6* cells (Figure S2). On the other hand, the CDK9 inhibitor Flavopiridol, which also blocks transcriptional elongation, but by preventing RNAPII-CTD-Ser2 phosphorylation(Bensaude, 2011), nearly eliminated *Mef2*+ nuclear dots (Figure S2). In either case, chemical inhibitor treatments supported the interpretation that intronic probe-positive nuclear dots represent nascent transcripts, which is indicative of zygotic *Mef2* expression in the PGCs during the tailbud developmental period. Notably, the temporal profile of zygotic *Mef2* expression showed an onset that followed both B7.6 cell remodeling and the global activation of RNA Polymerase II. This is consistent with a causal chain of events whereby postplasmic RNA exclusion through cellular remodeling helped relieve the Pem-1 break on RNAPII activity, thus permitting subsequent zygotic gene expression in the PGCs.

### JAK signaling delays the onset of zygotic *Mef2* expression in B7.6* cells

Having established that cellular remodeling precedes global RNAPII licensing and the onset of *Mef2* transcription in B7.6* cells, we sought to identify regulators of zygotic *Mef2* expression in the PGCs. We reasoned that signaling inputs from surrounding cells probably contribute to initiating zygotic gene expression in B7.6* cells, and their possible roles are readily testable by pharmacological inhibition. Treatments with U0126 (10μM; 10-12hpf; St. 21-23), Dorsomorphin (10μM; 10-12hpf) and 1-Azakenpaullone (10μM; 10-12hpf), which inhibit MEK1/2, BMPRI and GSK3, respectively, significantly reduced the proportions of nuclei with detectable nascent *Mef2* transcripts in *Ms4a15/2+* PGCs (Figure S3). However, neither of these inhibitors completely abolished zygotic *Mef2* expression. We thus tested whether these signaling pathways act additively, by combining the three inhibitors in a condition referred to as “3i” (10μM each; 10-12hpf), which indeed caused the most marked decrease in zygotic *Mef2* expression in B7.6* cells (Figure 3A-B, E). This suggested that MEK, BMPR and GSK3, whose possible sources are in the vicinity of PGCs at the tail tip and ventral side of the tail(Imai, 2004; Waki *et al*, 2015; Feinberg *et al*, 2019; Harder *et al*, 2019), act at least partially additively to promote zygotic *Mef2* expression in the PGCs.

**Figure 3.**
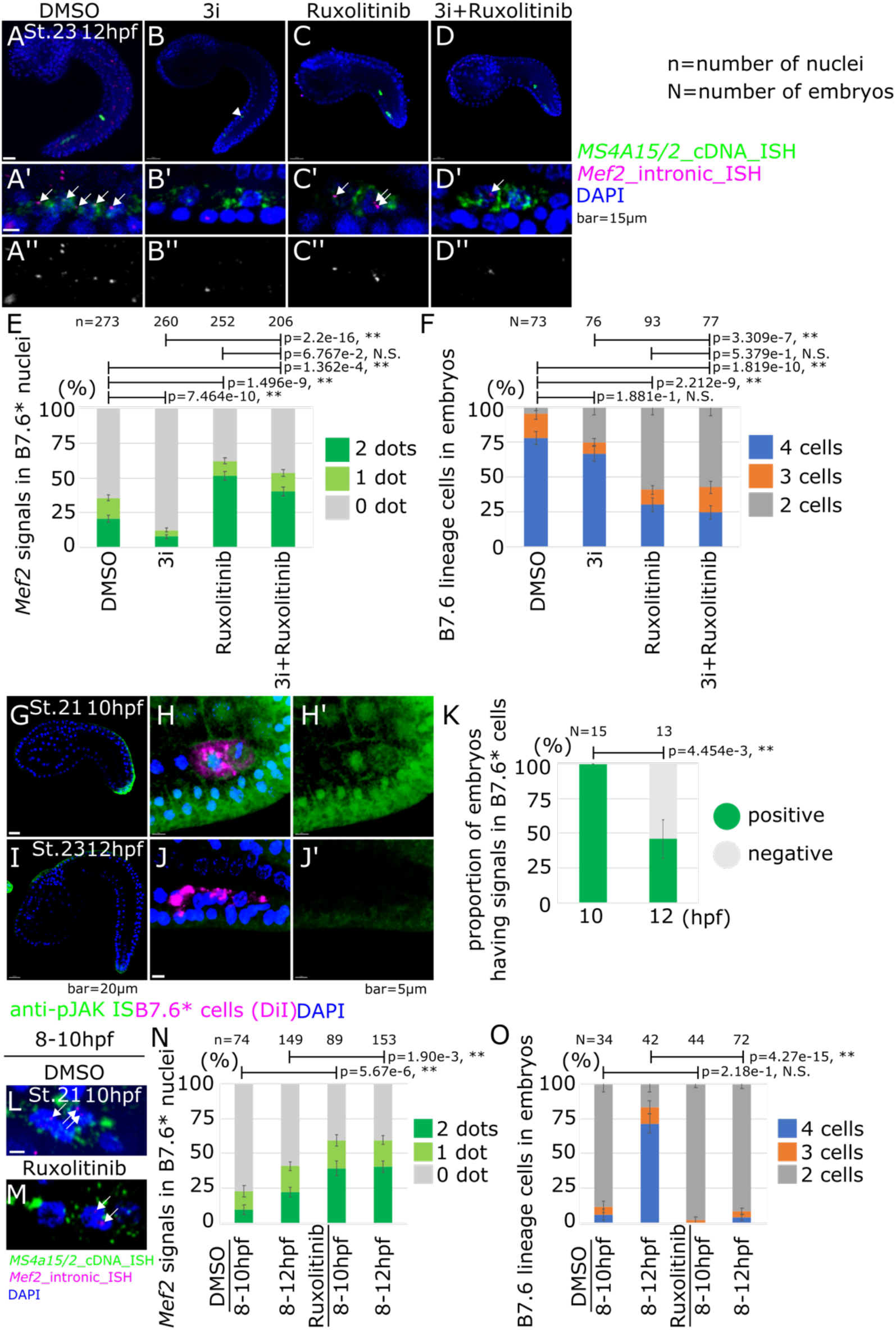
*Mef2* nascent expression in inhibitor treated embryos. (**A-D**) ISH was done with *Mef2* intronic probes under each pharmacological inhibitor treatment; DMSO (**A**), 3i (**B**), Ruxolitinib (**C**) and 3i+Ruxolitinib (**D**). White arrows show the dotted signals of nascent *Mef2* expression. White arrowhead shows lobe in (B). Scale bars=15μm. (A’-D’) Magnification images of A-D. (A’’-D’’) Black and white images of the magenta channel of A’-D’, respectively. (E) Proportion of *Mef2* signals in nuclei under each pharmacological inhibitor treatment. (F) Proportion of cell numbers of B7.6* cells in embryos under each pharmacological inhibitor treatment. (G-J) Immunostaining with anti-JAK2 antibody was done at the mid tailbud; 10 hpf (G-H’) and the late tailbud; 12 hpf (I-J’) stages. Scale bar=5μm. (K) Proportion of embryo that have positive signals in B7.6* cells by immunostaining with anti-JAK2 antibody. (L-M) ISH was done with *Mef2* intronic probes under each pharmacological inhibitor treatment; DMSO (L) and Ruxolitinib (M) in 10 hpf embryos. White arrows show the dotted signals of nascent *Mef2* expression. Scale bar=5μm. (N) Proportion of *Mef2* signals in nuclei under each condition. (O) Proportion of cell numbers of B7.6* cells in embryos under each condition. Error bars indicate standard error. p-value was calculated by z-test. p>0.05; N.S, 0.05>p>0.01; *, 0.01>p; **.

By contrast with the 3i treatment, the established JAK inhibitor Ruxolitinib (10μM; 10-12hpf) significantly increased the fraction of PGCs with *Mef2+* nuclei, in addition to inhibiting B7.6* cell division and tail elongation (Figure 3C, E-F). To test a possible hierarchy between the signaling pathways that regulate *Mef2* transcription in the PGCs, we combined the 3i and Ruxolitinib treatments (Figure 3D-E). This combination also increased the proportions of the *Mef2*+ nuclei, to levels similar to those obtained with Ruxolitinib alone. Taken together, these results suggest that JAK signaling inhibits zygotic *Mef2* expression in B7.6* cells, while MEK, BMPR and GSK3 act additively to promote *Mef2* transcription, antagonizing JAK activity.

To gain further insights into the biological significance of inhibitor treatments and better characterize the context in which JAK signaling regulates zygotic *Mef2* expression in the PGCs, we sought to assay endogenous JAK activity in *Ciona* embryos. Previous genome-wide surveys identified two *Jak* ortholog (*Jak-a*: KH.C1.555 and *Jak-b*: KH.C8.409) in the *Ciona* genome(Hino *et al*, 2003; Tokuoka *et al*, 2022). The *Jak-a* RNAs are maternally deposited in unfertilized eggs as detected by ISH, ESTs and bulk RNA-seq (Supplementary file 2)(Satou *et al*, 2005),(Dardaillon *et al*, 2020). *Ciona* Jak-a displays one region surrounding a potential active phosphorylated tyrosine residue that is conserved with human JAK2 (Figure S4A). We thus tested an anti phospho-human JAK2 (Y931) polyclonal antibody to detect endogenous JAK activity in tailbud embryos, and observed conspicuous signal in tail tip and dorsal midline cells (Figure 3G, I). Treatment with the JAK inhibitor Ruxolitinib abolished the signal, which is consistent with specificity of both the drug treatment and antibody staining (Figure S4B-G). JAK activity was most conspicuous at the tail tip of St. 21 embryos (10 hpf at 18°C), and markedly reduced by St. 23 (12 hpf at 18°C), which coincides with peak transcriptional activity of the *Mef2* gene in the germline at St. 23 (Figure 2F). While this pattern is consistent with concomitant downregulation of JAK signaling and activation of *Mef2* expression, as well as the effect of Ruxolitinib treatments, the latter do not distinguish between cell autonomous and non cell autonomous effects in B7.6* cells. To evaluate the possibility that JAK acts cell-autonomously upstream of *Mef2* to repress its transcription in the PGC lineage, we DiI-labeled B7.6 cells and assayed JAK activity at two developmental stages corresponding to the onset and peak of zygotic *Mef2* expression (Figure 3G-K). This experiment revealed JAK activity in B7.6* cells at St. 21 (10 hpf at 18°C), but reduced signaling by St. 23 (12 hpf at 18°C), which is consistent with a cell-autonomous inhibitory effect on early *Mef2* transcription, although it does not formally rule out non-cell autonomous influence from the distal tail epidermis.

Of note, the localization of maternal *Jak-a* mRNAs in early embryos, opens the possibility that JAK signaling is active in early B7.6 blastomeres. We tested endogenous JAK activity at early developmental stage, and observed signals at vegetal side of cell membrane on B7.6 cells and endodermal cells (Figure S5A). JAK activity initiates during the 110-cell to early gastrulation stage (Figure S5B-C). Treatment with a translation-inhibitor antisense morpholino oligonucleotide conjugated vivo (vivo-MO) targeting the 5’ untranslated region (5’ UTR) of *Jak-a* mRNA abolished the anti-pJAK2 signal, indicating that maternal Jak-a mediates JAK signaling, and further supporting the notion that both the vivo-MO treatment and antibody staining are specific tools to study JAK signaling in *Ciona* (Figure S5C-D). In summary, these data indicate that maternal *Jak-a* mRNA fosters JAK signaling in the germline, from early gastrulation and until the tailbud stage.

Leveraging anti-pJAK immunostaining, we tested whether MEK, BMPR and GSK3 signaling contribute to inhibiting JAK phosphorylation by stage 23 by staining embryos treated with the 3i cocktail of inhibitors (Figure S6). We observed no conspicuous difference between 3i and DMSO control treatments, suggesting that Jak dephosphorylation occurs independently of MEK, BMPRI, GSK3 signaling inputs (Figure S6).

Taken together, our observations are consistent with the possibility that JAK signaling plays an early inhibitory role in delaying the peak of *Mef2* transcription in the PGCs. Indeed, RNA polymerase II is licensed to initiate transcription as early as 8 hpf (Figure 2D; St. 16), while zygotic *Mef2* expression does not start before 10 hpf, and only peaks at 12 hpf (Figure 2F). To test if JAK signaling inhibits early *Mef2* expression, we thus treated embryos with Ruxolitinib from 8 hpf onward (St. 16), and assays zygotic *Mef2* expression at 10 hpf (Figure 3L-O). In control DMSO-treated embryos, at most 25% of the B7.6 lineage nuclei showed nascent *Mef2* transcripts. By contrast, we detected active *Mef2* transcription in approximately 50% of the B7.6 lineage nuclei following treatment with the JAK inhibitor between 8 and 10 hpf (St. 16-21; Figure 3L-O). Moreover, 8 to 12 hpf (St. 16-23) treatments resulted in similar ~50% of *Mef2^intron^+* B7.6 lineage nuclei, albeit with a lower fold-increase, as the DMSO baseline was at 41%, suggesting that *Mef2* transcription peaks earlier following early JAK inhibition. Of note, control B7.6* cells had not divided in the 8 to 10 hpf time window (Figure 3O). These early treatments thus indicated that chemical JAK inhibition promotes *Mef2* transcription independently of its effects on cell division. Taken together, these results support the notion that early, possibly cell-autonomous, JAK activity delays the peak, and probably the onset of zygotic *Mef2* expression in the PGCs.

### MEK, BMPR, GSK3 and JAK signaling regulate *Mef2* transcription independently of RNA polymerase II CTD phosphorylation

Finally, since global transcription licensing shortly precedes the onset of zygotic *Mef2* expression, we tested whether the kinases that influence the production of nascent *Mef2* transcripts also impact RNAPII-CTD phosphorylation. To this aim, we repeated the above chemical inhibitor treatments targeting MEK, BMPRI, GSK3 and JAK, and assayed RNAPII-CTD-Ser2 phosphorylation at 8 hpf, St. 16 and 12 hpf, St. 23. In short, neither of these inhibitor treatments appeared to alter RNA polymerase II activity (Figure 4A-E; Figure S7), suggesting that the kinases regulate zygotic *Mef2* expression independently of global RNAPII activation (Figure 4G).

**Figure 4.**
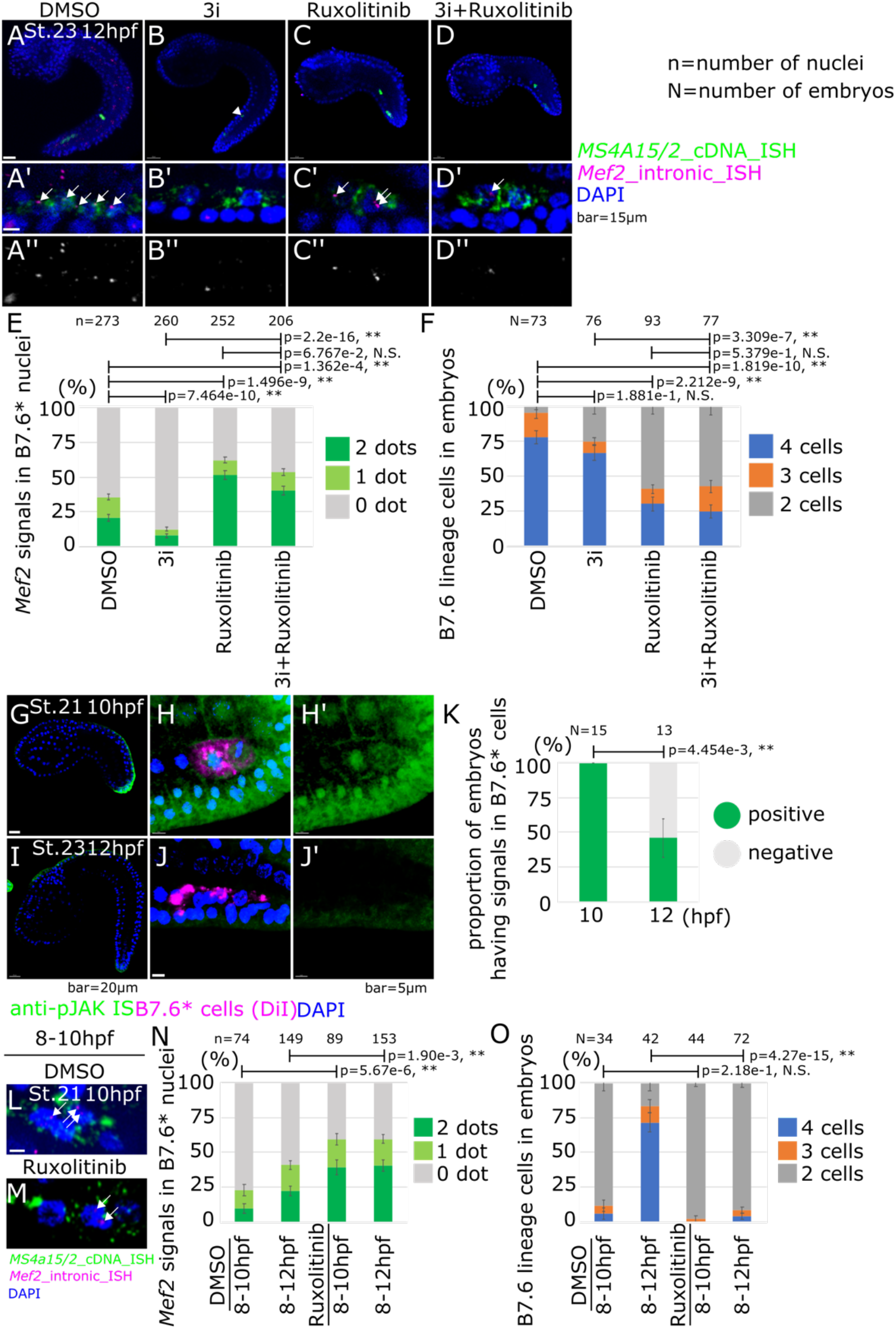
RNA polymerase II Ser2 phosphorylation in inhibitor treated embryos. (A-C) Immunostaining was done with anti-RNAPII-CTD-pSer2 antibody under each pharmacological inhibitor treatment at 12 hpf. Scale bar=5μm. (D) Proportion of signals of immunostaining with anti-RNAPII-CTD-pSer2 antibody in nuclei under each pharmacological inhibitor treatment. (E) Proportion of cell numbers of B7.6* cells in embryos under each pharmacological inhibitor treatment. Error bars indicate standard error. p-value was calculated by z-test. p>0.05; N.S, 0.05>p>0.01; *, 0.01>p; **. (F) A schematic image for activity or presence of each factor in B7.6 and B7.6* cells based on this study and the previous report (Shirae-Kurabayashi *et al*, 2011). (G) Working scheme of onset of zygotic transcription in B7.6* cells that was drawn in this study.

## Discussion

Animal species endowed with “preformed’’ primordial germ cells illustrate August Weismann’s famous dichotomy between mortal somatic vs. immortal germline lineages, and the mechanisms underlying this divergence in early embryos have garnered much attention(Kutschera & Niklas, 2004; Dröscher, 2014; Weismann, 1892). Despite the *de facto* immortality and totipotency of their lineage, illustrated by their ability to reconstitute an entire organism upon fertilization and embryogenesis, PGCs eventually assume an identity and differentiate into highly specialized egg and sperm cells, the gametes. As cells, PGCs are thus bound to the same developmental constraints as somatic cells: to choose an identity, which comprises avoiding others, and differentiate accordingly, with the notable distinction that germ cells may postpone terminal differentiation until sexual maturity(Kang & Han, 2011; Hayashi & Saitou, 2014; Bhartiya *et al*, 2017).

This opportunity to pause differentiation may relate to the observed delay in starting zygotic gene expression in the germline, which is a widespread mechanism used to prevent somatic fate specification(Strome & Updike, 2015). Indeed, the unrelated proteins Pgc, PIE1 and Pem-1 globally prevent zygotic gene expression in the germline by inhibiting phosphorylation of Serine 2 in heptapeptide repeats of the C-terminal domain of RNA Polymerase II, in as diverse animals as Drosophila, *Caenorhabditis elegans* and ascidians, respectively(Seydoux & Dunn, 1997; Batchelder *et al*, 1999; Blackwell, 2004; Seydoux & Braun, 2006; Hanyu-Nakamura *et al*, 2008).

This early global inhibition of transcription in PGCs imposes an exclusive reliance on maternal products deposited in the egg during oogenesis(Lehmann, 2012; Trcek & Lehmann, 2019). Among those, germline-specific components are typically found in RNA-rich germ granules, germ plasm or nuage, and dispatched into PGCs through polarized RNA and protein localization, and asymmetric inheritance following unequal cleavages in early embryos(Sardet *et al*, 2007; Little *et al*, 2015; Trcek & Lehmann, 2019). In early ascidian embryos, *Pem-1* maternal mRNAs are localized and associated with the cortical centrosome attracting body (CAB), segregate asymmetrically at every unequal cleavage, and are found exclusively in B7.6 blastomeres by the 64-cell stage(Sardet *et al*, 2007). While the Pem-1 protein keeps B7.6 cells, and their progenitors, transcriptionally quiescent, it does not have a direct role in specifying the germline identity, which is another function that must be partially carried by the germ plasm, in the absence of zygotic gene expression(Strome & Updike, 2015; Nakamura & Seydoux, 2008). In *Ciona*, as in numerous other species, the conserved RNA helicase Vasa/Ddx4 is thought to act as a PGC determinant(Takamura *et al*, 2002). Vasa/Ddx4 mRNAs and proteins are segregated mainly to B7.6* cells/PGCs, as is the case for P granules in *C. elegans*(Paix *et al*, 2009; Abdu *et al*, 2016). The germ plasm must thus fulfill two essentials, but partially antagonistic functions for germline specification: (1) inhibiting the alternative somatic fate specification through transcriptional silencing, and (2) promoting the acquisition of germ line-specific molecular features, including transcriptional activation of a zygotic germline program. We must thus anticipate that germ plasm remodeling and segregation of anti-somatic and pro-germline activities is essential for germline development. Consistent with this assertion, germ plasm remodeling has been observed, and segregation of its components is important for germline development in flies(Little *et al*, 2015) and zebrafish(D’Orazio *et al*, 2021).

Here, we presented data that corroborate previous observations about the dislocation of the germ plasm and differential segregation of its components in *Ciona*, and showed that this results from a peculiar cellular remodeling event rather than unequal cell cleavage, as previously thought based on phospho-histone 3 staining in B7.6(Shirae-Kurabayashi *et al*, 2006). The vesicles had thus been referred to as B8.11 cells, but did not show DAPI+ staining or evidence of nucleokinesis, and are actually not cells. We propose to call them “lobes”, after the analogous entity observed in *C. elegans*. Moreover, *Pem-1* mRNAs are among the postplasmic RNAs that segregate specifically to the lobe(Yamada, 2006; Paix *et al*, 2009), ultimately resulting in the elimination of Pem-1 proteins from the PGCs by the tailbud stage(Shirae-Kurabayashi *et al*, 2011). Similar patterns of postplasmic RNA segregations were observed in other tunicates, like the Appendicularian, *Oikopleura dioica*(Olsen *et al*, 2018). Since Pem-1 globally blocks transcription, we propose that lobe scission is an important step to physically remove postplasmic RNAs from the PGCs and permit further germline specification, in part through zygotic gene expression. In another ascidian species, *Halocynthia roretzi*, removal of *Pem-1* gene products appears to be necessary and sufficient for zygotic expression of ATP/ADP translocase gene in the germline; however, the cellular and molecular mechanisms likely differ as a sizeable portion of the *HrPEM* RNAs are retained in B7.6* PGCs(Miyaoku *et al*, 2018).

Germline lobe formation and scission exhibit intriguing parallels and differences between *Ciona* and *C. elegans*. One of the hallmarks of lobe formation and scission in *C. elegans* is the polarized distribution and removal of most mitochondria from the cell bodies of PGCs(Abdu *et al*, 2016; Schwartz *et al*, 2022). By contrast, in *Ciona* embryos, most mitochondria are depleted from B7.6 cells during the asymmetric division of their mother B6.3 cell, which also produces the B7.5 cardiac progenitors that inherit most mitochondria (a.k.a. myoplasm in ascidians)(Chenevert *et al*, 2013). In *Ciona*, mitochondria clearance from the germline is thus decoupled from lobe formation.

Another intriguing parallel between germline lobe formation and scission in *C. elegans* and *Ciona* regards its association with endodermal progenitors. Primordial germ cells exhibit an evolutionary conserved association with the endoderm in a variety of species(Anderson *et al*, 2000; Kanamori *et al*, 2019), including *Ciona*(Karaiskou *et al*, 2015; Kawai *et al*, 2015; Krasovec *et al*, 2019). The physical proximity between lobes and endoderm cells could thus be shared between *Ciona* and *C. elegans* as a byproduct of an ancient association between PGCs and endodermal progenitors. As lobe formation in *C. elegans* is inhibited by loss of dynamin and Rho function in the endoderm(Abdu *et al*, 2016; Maniscalco *et al*, 2020), germline lobes processing in *Ciona* could also involve the endoderm. Perturbation of endodermal functions would be required to test this hypothesis, and may reveal further cellular differences in the relationships between the endoderm and the germline in *Ciona* and *C. elegans*.

Finally, much attention has been devoted to the molecular mechanisms underlying germ plasm structure and function during germline formation(Seydoux & Braun, 2006; Lehmann, 2012; Trcek & Lehmann, 2019). By contrast, less is known about the mechanisms involving cell-cell signaling and zygotic gene expression in early germline specification, with the notable exception of PGC specification in mammals, which is governed by FGF, BMP and Wnt signaling pathways, as well as leukemia inhibitory factor (LIF)-controlled signaling(Lawson *et al*, 1999; Ohinata *et al*, 2009; Yu *et al*, 2020). Here, having identified one zygotically expressed gene in *Ciona* PGCs, we began to disentangle the mechanisms governing early genome activation in the germline. Specifically, chemical inhibitor treatments suggest that MEK, BMPRI and GSK3 contribute to activating *Mef2* transcription in the PGCs, providing JAK activity is inhibited. Our observations indicate that early JAK signaling acts independently of Pem-1-mediated inhibition of RNAPII to inhibit *Mef2* transcription, and after lobe scission to delay the onset of *Mef2* expression. Whereas, further studies will be required to dissect cell autonomous or non cell autonomous effects in B7.6* cells, and to determine the source of signals and the genome-wide extent of zygotic gene expression in the *Ciona* germline, these results evoke a parallel between the signaling circuits that control germline specification in *Ciona*, and those that maintain stemness in mammalian pluripotent stem cells, which also rely on active JAK signaling and the concurrent inhibition of MEK and GSK3(Ying *et al*, 2008; Hanna *et al*, 2010; Gafni *et al*, 2013; Ware *et al*, 2014; Takashima *et al*, 2015).

## Materials and Methods

### Animals

Wild-type animals of *Ciona intestinalis* type A were collected by M-Rep (Marine Research and Educational Products’), in San Diego, CA, and *Ciona intestinalis* type B were collected in Bergen, Norway. While *Ciona intestinalis* type A and type B have recently been considered as different species(Brunetti *et al*, 2015), they can cross each other to establish hybrid animals, who can be raised to at least F3 generation, suggesting their conserved developmental process(Suzuki *et al*, 2005; Sato *et al*, 2014; Malfant *et al*, 2017; Ohta *et al*, 2020). In this study, we used both *Ciona* species in mixture. Especially, data in Supplemental Figure 1 and 5 were taken in *Ciona intestinalis* type B, and the others were taken in *Ciona intestinalis* type A. Eggs and sperm were surgically collected from mature adults. Chorion of fertilized eggs were removed by Sodium thioglycolate and Proteinase as previously described(Christiaen *et al*, 2009a). Dechorionated eggs were cultured on agarose coated Petri dishes in TAPS-buffered artificial sea water (ASW; Bio actif sea salt, Tropic Marin).

### DiI cell tracing

CellTracker CM-DiI Dye (Thermo Fisher Scientific) was dissolved in DMSO (Fisher Scientific) to 1 mg/mL as previously reported(Satou *et al*, 2004). The DiI solution was sprayed with a microneedle onto B7.6 cells of 64-cell stage embryos. These embryos were allowed to develop, fixed with MEM-FA or MEM-PFA (3.7% Formaldehyde or 4% Paraformaldehyde, 0.1M MOPS, 0.5M NaCl, 1mM EGTA, 2mM MgSO_4_), and used for antibody staining and fluorescent *in situ* hybridization (FISH).

### Antibody staining

We used an antibody for RNA polymerase II as a previous report(Shirae-Kurabayashi *et al*, 2011); CTD-pSer2 (Abcam, ab5095, 1:500 dilution). We followed the previously described protocol(Ohta & Satou, 2013) with slight modification. A rabbit anti human phospho-JAK2 (Y931) antibody (Thermo Fisher Scientific, PA5-104704) was used as a primary antibody, 1/500 in *Can Get Signal* Immunostain Solution A (TOYOBO). The antibody was detected by an anti-rabbit-HRP goat antibody 1/500 in Can Get Signal Immunostain Solution A (TOYOBO), and by Tyramide Signal Amplification (Perkin Elmer) as previously described(Ohta & Satou, 2013).

### *In situ* hybridization

DNA fragments were amplified by PCR with exTaq-HS (Takara Bio) and Phusion HF (New England Biolabs) DNA polymerases from *Ciona* genomic DNA or cDNA. The primers that we used were summarized in supplementary file 1. The amplicons were subcloned into TOPO vectors (life technologies). DIG or fluorescein labeled RNA probes were synthesized by T7 and sp6 RNA polymerases (Roche) from template DNA plasmid digested by NotI or SpeI (New England Biolabs), and were cleaned by RNeasy mini kit (QIAGEN). We followed the protocol for *in situ* hybridization described before(Christiaen *et al*, 2009b; Ohta & Satou, 2013). We detected fluorescein and DIG probes using TSA plus (Perkin Elmer) green (FP1168) and red (FP1170), respectively. Primer sequences are provided in supplementary file 1.

We used an antibody for phospho histone 3 (pH3) that was previously reported(Shirae-Kurabayashi *et al*, 2006) (pH3-ser10-6g3-mouse-mAb, Cell Signaling; #9706; 1:500 diluted). The primary antibody was added together with anti-DIG antibody during FISH process, and secondary antibody (anti-mouse-Alexa-555, Thermo Scientific, A-21127) was used to detect the anti-pH3 antibody after detection of ISH probe by using TSA plus (Perkin Elmer) green (FP1168).

### Pharmacological inhibitor treatments

Actinomycin D (50-76-0; A1410; Sigma) was diluted into DMSO at 10 mg/mL stock. The stock solution was diluted into ASW to a final concentration of 40 μg/mL. This concentration was reported to block transcription in *Halocynthia* embryos(Miyaoku *et al*, 2018). Flavopiridol (146426-40-6; S1230; Selleck chemicals) was diluted into water to 10mM stock. The stock solution was diluted into ASW to final concentration 1 and 10 μM. The transcriptional inhibitor treated embryos were fixed by MEM-PFA (4% PFA, 0.1M MOPS, 0.5M NaCl, 1mM EGTA, 2mM MgSO_4_) after 1 hour inhibitor treatment, and used for *in situ* hybridization.

1-Azakenpaullone (S7193; Selleckchem; (Feinberg *et al*, 2019)), Ruxolitinib (INCB018424; S1378; Selleckchem), Vismodegib (GDC-0449; S1082; Selleckchem), DAPT (208255-80-5; D5942; Millipore Sigma), SB431542 (S1067; Selleckchem; (Ohta & Satou, 2013)), U0126 (9903; Cell Signaling Technology; (Hudson *et al*, 2003)) and Dorsomorphin (1219168-18-9; S7306;Selleckchem; (Ohta & Satou, 2013; Feinberg *et al*, 2019)) were used to perturb define signaling pathways as described in corresponding references. These treatments were done in a final concentration of 10 μM for 2 or 4 hours. The inhibitor-treated embryos were fixed by MEM-PFA after 2 hours inhibitor treatment, and used for *in situ* hybridization.

### Vivo-morpholino oligonucleotide treatment

We designed antisense morpho oligonucleotide conjugated vivo (vivo-MO) at the 5’ utr of *Jak-a* gene (5’-CTTTTGGTTAGCATGAATTGAAGCC-3’). The stock solution of vivo-MO was made by diluting with water to be 0.5 mM. Dechorionated eggs were treated with 20 μM, 40 μM and 60 μM for 20 minutes during 25 to 45 minutes after fertilization in 250 μL ASW in 2mL microcentrifuge tube, and transferred into ASW on agarose coated Petri dishes until fixation with MEM-FA. Mock control embryos were treated with water the same volume of that of 60 μM vivo-MO condition. The fixed embryos were used for immunostaining. Because the previous report used vivo-MO in Zebrafish eggs and showed that 40 μM and 60 μM concentration of vivo-MO worked to block function of target gene(Wong & Zohar, 2015), we tested the same concentration here.

We used representative data on the figures after reproducing results in at least two independent experiments.

## Acknowledgments

We thank Pr. Hiroki Nishida for helpful comments on the original version of this manuscript. We are grateful to Christiaen lab members for discussions and feedbacks. This work was supported by NIH/NIGMS award GM096032 to L.C.

---

**Supplemental Figure 1.**
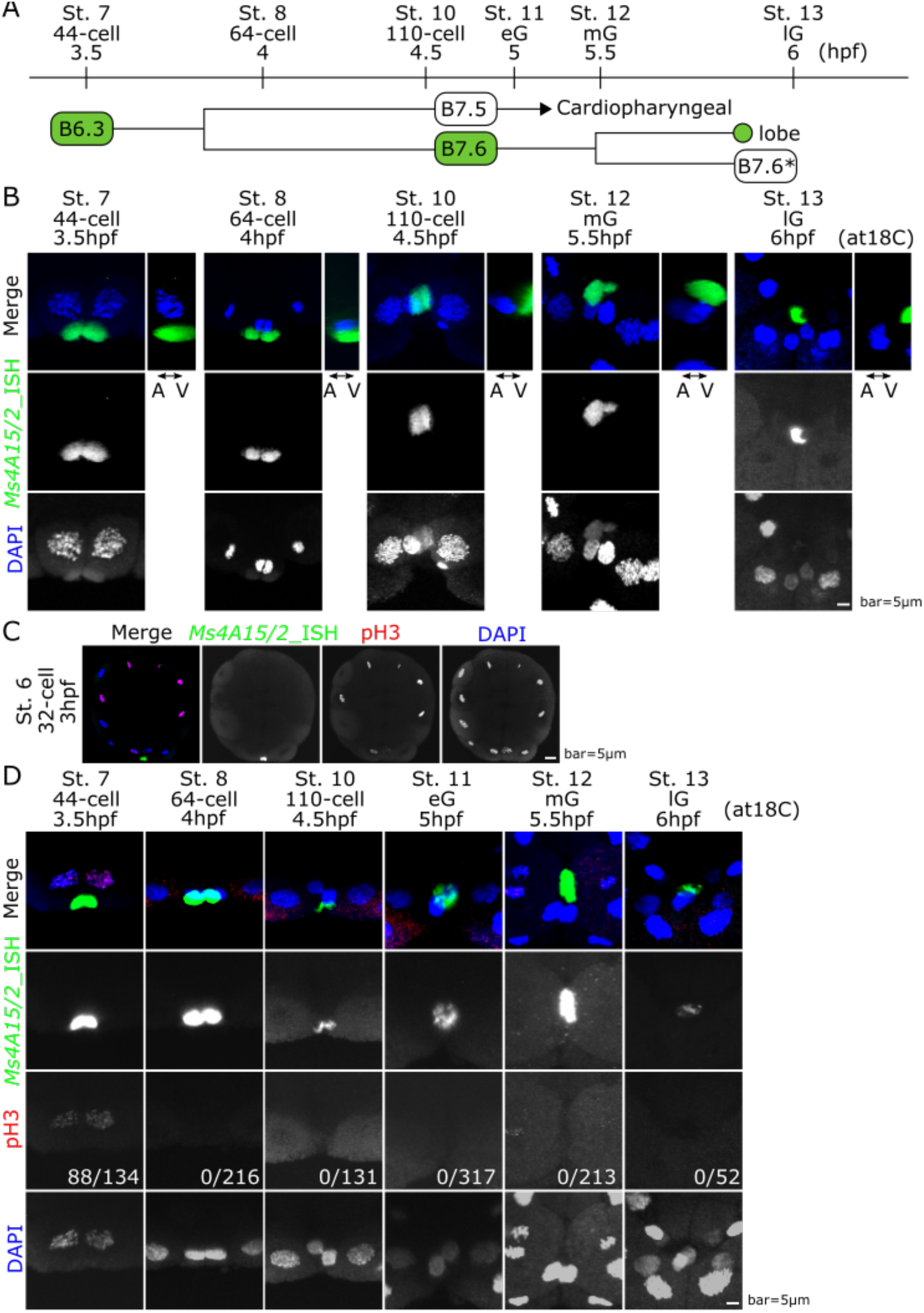
(A) Time scale of cell division from 32-cell stage to late gastrula stage in *Ciona* embryo. (B) Time series embryos from 44-cell stage to late gastrulation (lG) stage were stained by PGC marker *Ms4A15/2* by FISH. These pictures are vegetal view. The pictures on the side are lateral view. (C) The embryos at 32-cell stage were stained by both PGC marker *Ms4A15/2* by FISH and anti-pH3 antibody. (D) Time series embryos from 44-cell stage to late gastrulation (lG) stage were stained by both PGC marker *Ms4A15/2* by FISH and anti-pH3 antibody. Numbers shows the pH3 signal positive in B6.3 or B7.6 cells out of total embryos. These pictures are vegetal view. Nuclei were stained with DAPI showed in blue.

**Supplemental Figure 2.**
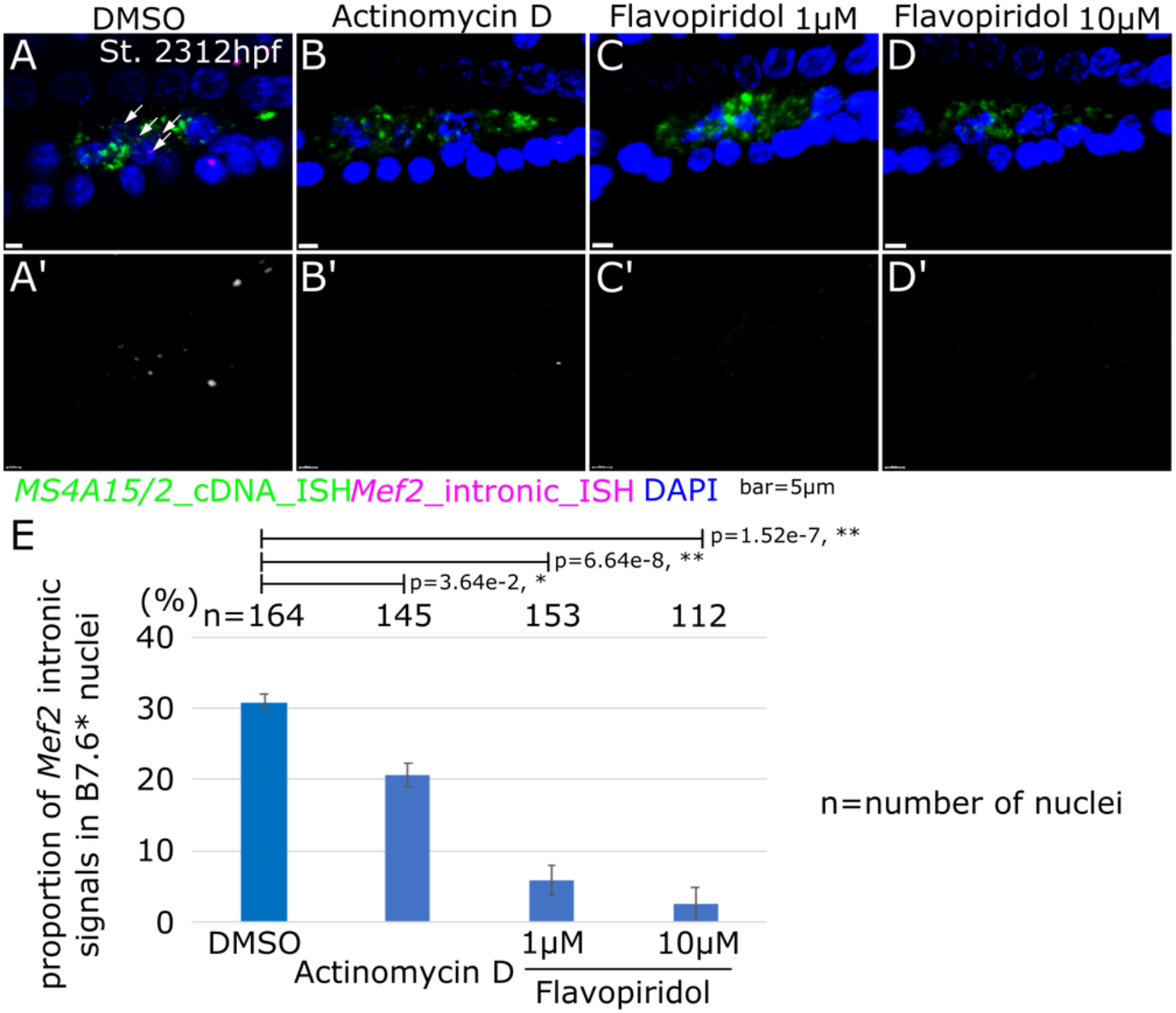
(A-D’) Nascent expression of *Mef2* in embryos treated by: DMSO (A), Actinomycin D (B), 1μM (C) and 10μM (D) Flavopiridol. A’-D’ shows black and white image of the magenta channel. White arrows indicate the dotted signals of nascent *Mef2* expression in nuclei of B7.6* cells. Scale bar=5μm (E) Proportion of *Mef2* signals in nuclei in B7.6* cells. Error bars indicate standard error. p-value was calculated by z-test. p>0.05; N.S, 0.05>p>0.01; *, 0.01>p; **.

**Supplemental Figure 3.**
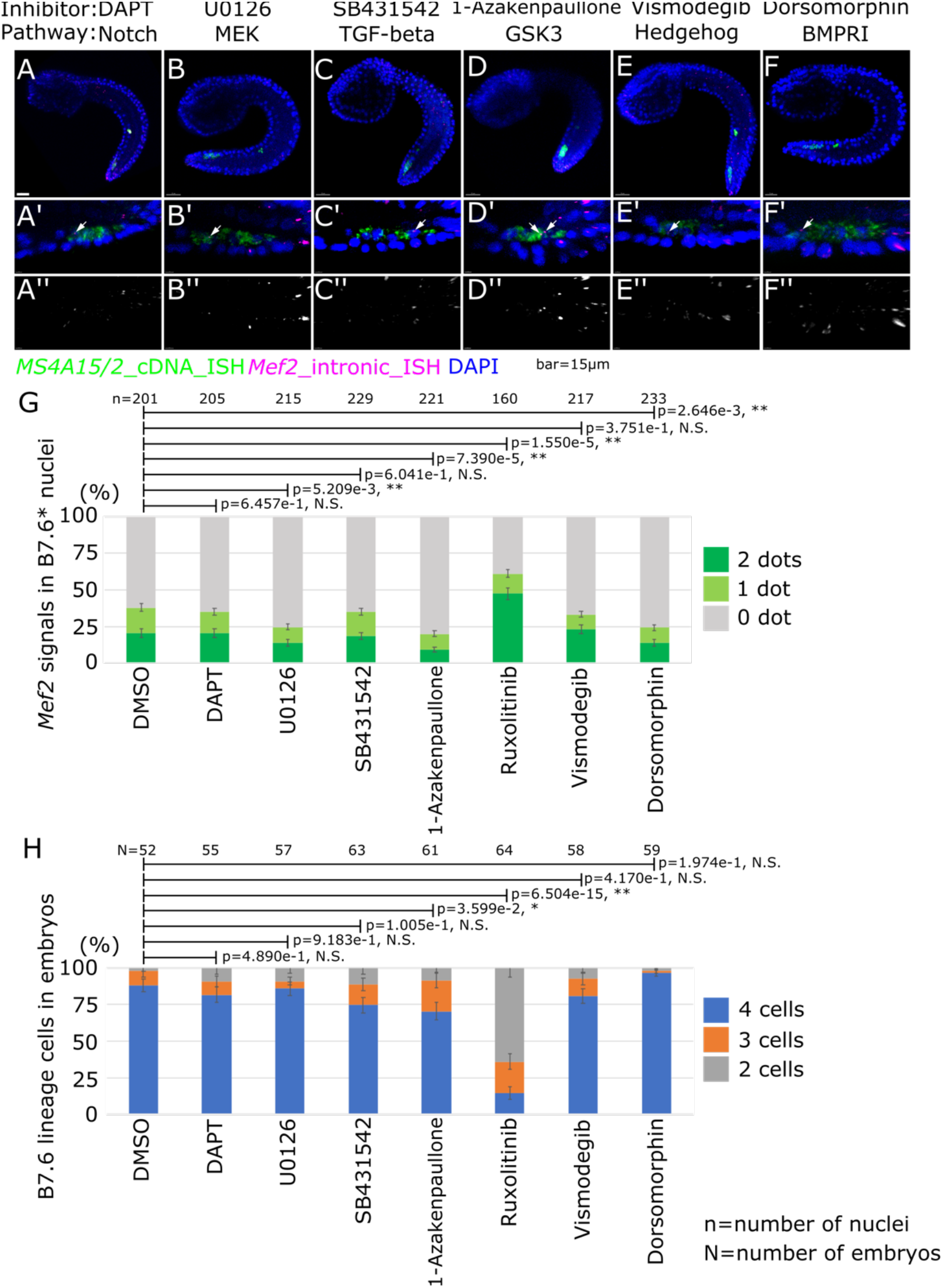
(A-F) ISH was done with *Mef2* intronic probes under each pharmacological inhibitor treatment. White arrows indicate the dotted signals of nascent *Mef2* expression in nuclei of B7.6* cells. Scale bar=15μm. (A’-F’) Magnification images around B7.6* cells. (A’’-F’’) Black and white image of the magenta channel of A’-F’, respectively. (G) Proportion of *Mef2* signals in nuclei of B7.6* cells under each pharmacological inhibitor treatment. (H) Proportion of cell numbers of B7.6* cells in embryos under each pharmacological inhibitor treatment. Error bars indicate standard error. p-value was calculated by z-test. p>0.05; N.S, 0.05>p>0.01; *, 0.01>p; **.

**Supplemental Figure 4.**
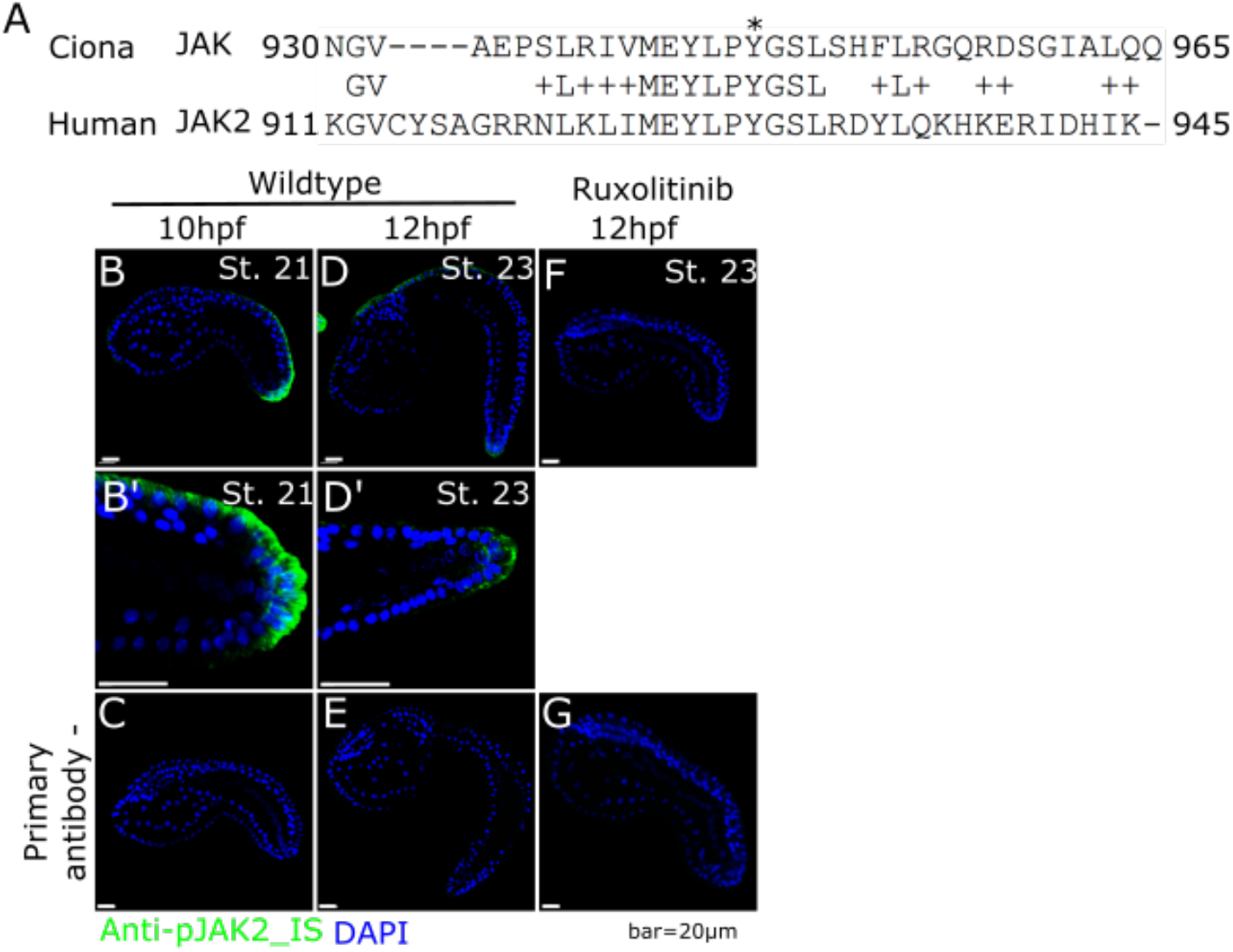
(A) Aliment of protein sequences of JAK in *Ciona* and JAK2 in Human. Amino acid residues 930-965 in *Ciona* JAK are shown. The conserved actively phosphorylated Tyrosines is marked by *. (B, D, F) Immunostaining was done with anti-pJAK2 antibody in DMSO control at 10 hpf (B) and 12 hpf (D), and Ruxolitinib treated embryos at 12 hpf (F). (B’, D’) Images around the tail tip of B and D, respectively. (C, E, G) No primary antibody controls corresponding to B, D and F, respectively. Scale bar=20μm.

**Supplemental Figure 5.**
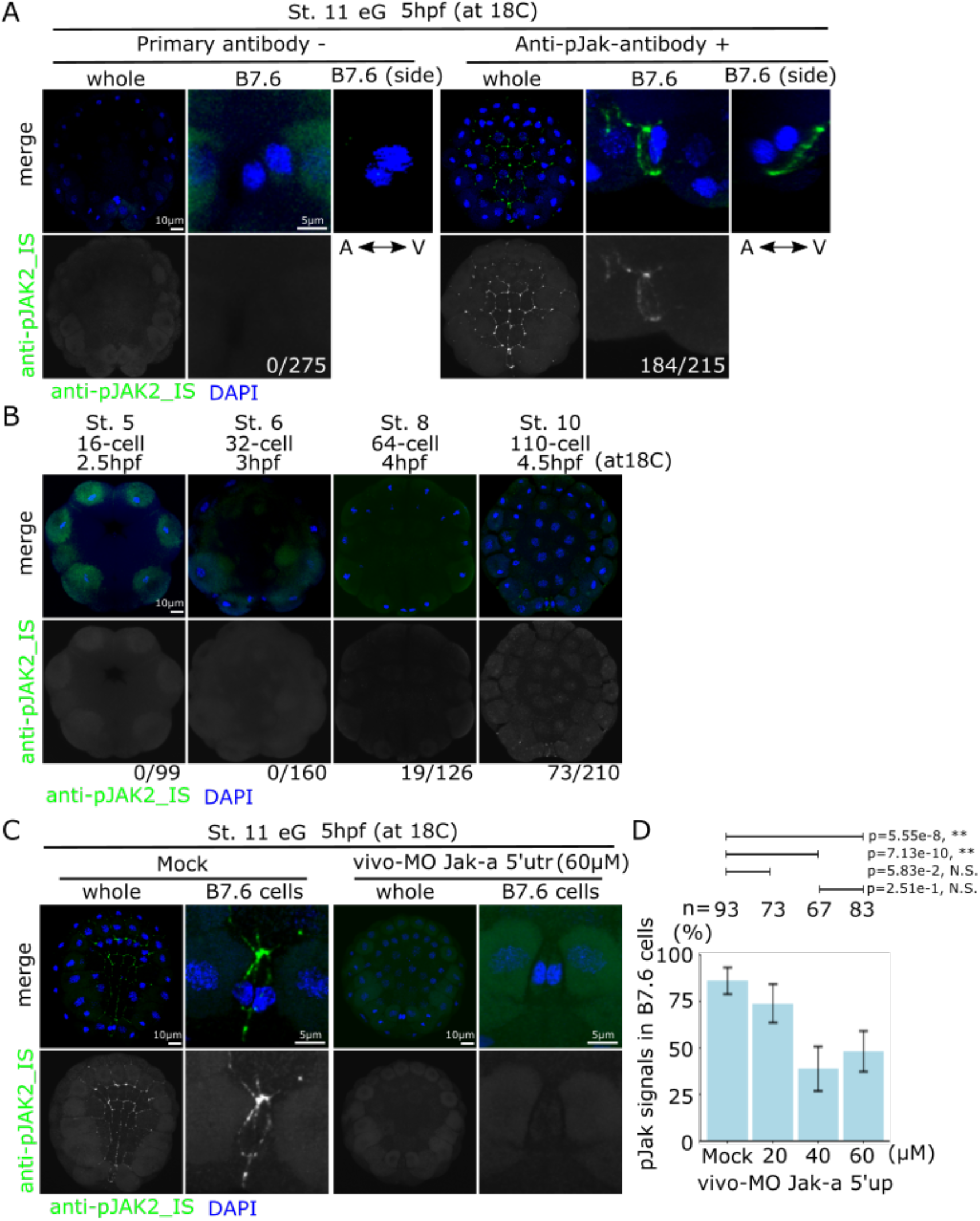
(A) Immunostaining was done with anti-pJAK2 antibody at early gastrulation (eG) stage with no primary antibody control. The numbers show the embryo showing signals on B7.6 cells out of total embryo. These pictures of whole embryo (whole) and B7.6 cells (B7.6) were taken from vegetal side. The pictures on the side were taken from the side of B7.6 cells. (B) Time series embryos from 16-cell stage to 110-cell stage were stained with anti-pJAK2 antibody immunostaining. These pictures of whole embryo were taken from vegetal side. The numbers show the embryo showing signals on B7.6 cells out of total embryo. (C) The dechorionated eggs were treated in water (Mock) or vivo-MO for *Jak-a* 5’ UTR and performed immunostaining with anti-pJAK2 antibody. (D) Rate of embryos showing signals on B7.6 cells. n means the number of embryos. p-value was calculated by t-test. p>0.05; N.S, 0.05>p>0.01; *, 0.01>p; **. Nuclei were stained with DAPI showed in blue. Scale bar=10μm and 5μm.

**Supplemental Figure 6.**
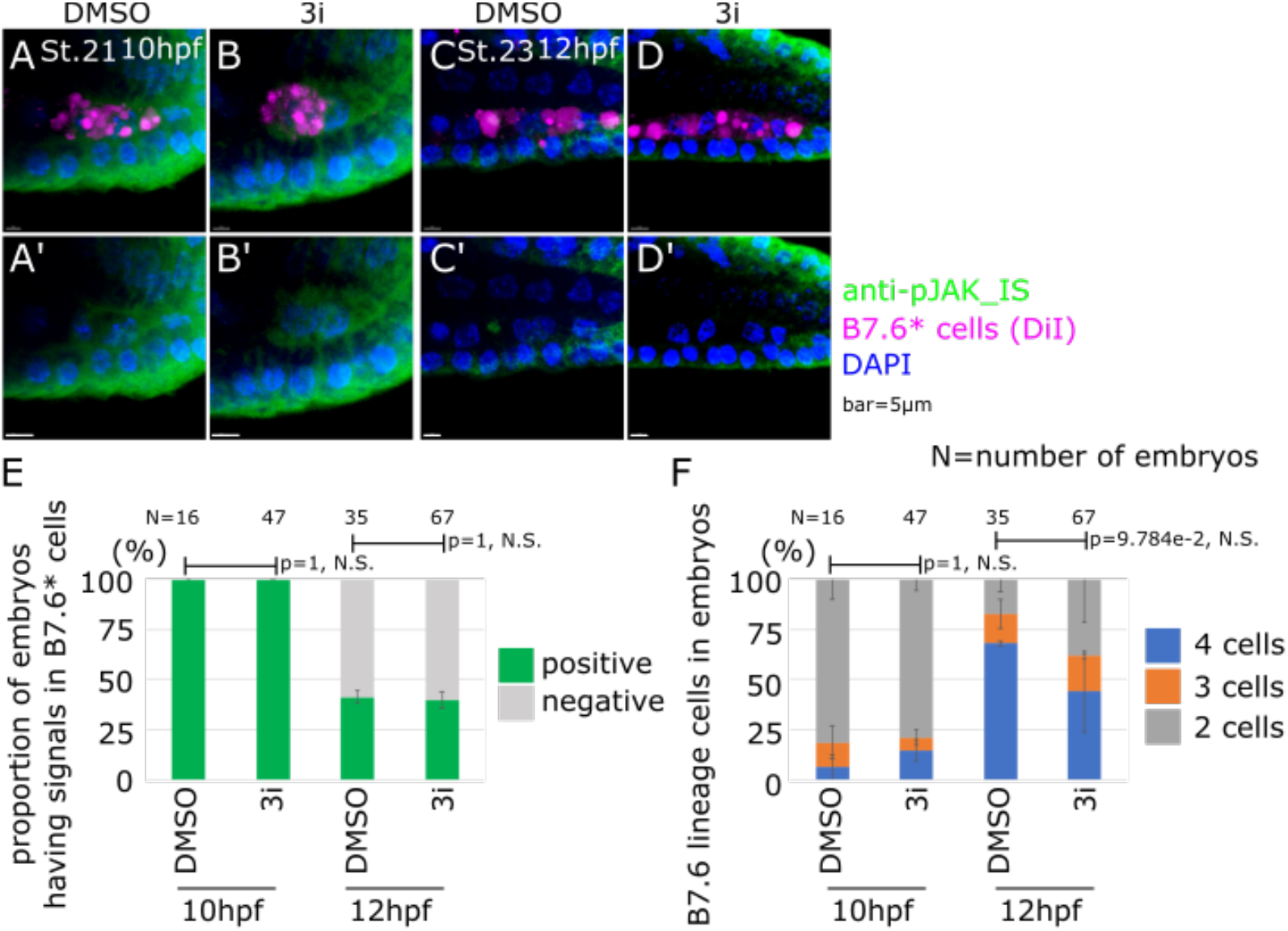
(A-D) Immunostaining was done with anti-JAK2 antibody under 3i pharmacological inhibitors treatment at 10hpf (A, B) and 12hpf (C, D). Scale bar=5μm. (E) Proportion of signals of immunostaining with anti-pJAK2 antibody in 7.6* cells. (F) Proportion of cell numbers of B7.6* cells in embryos. Error bars indicate standard error. p-value was calculated by z-test. p>0.05; N.S, 0.05>p>0.01; *, 0.01>p; **.

**Supplemental Figure 7.**
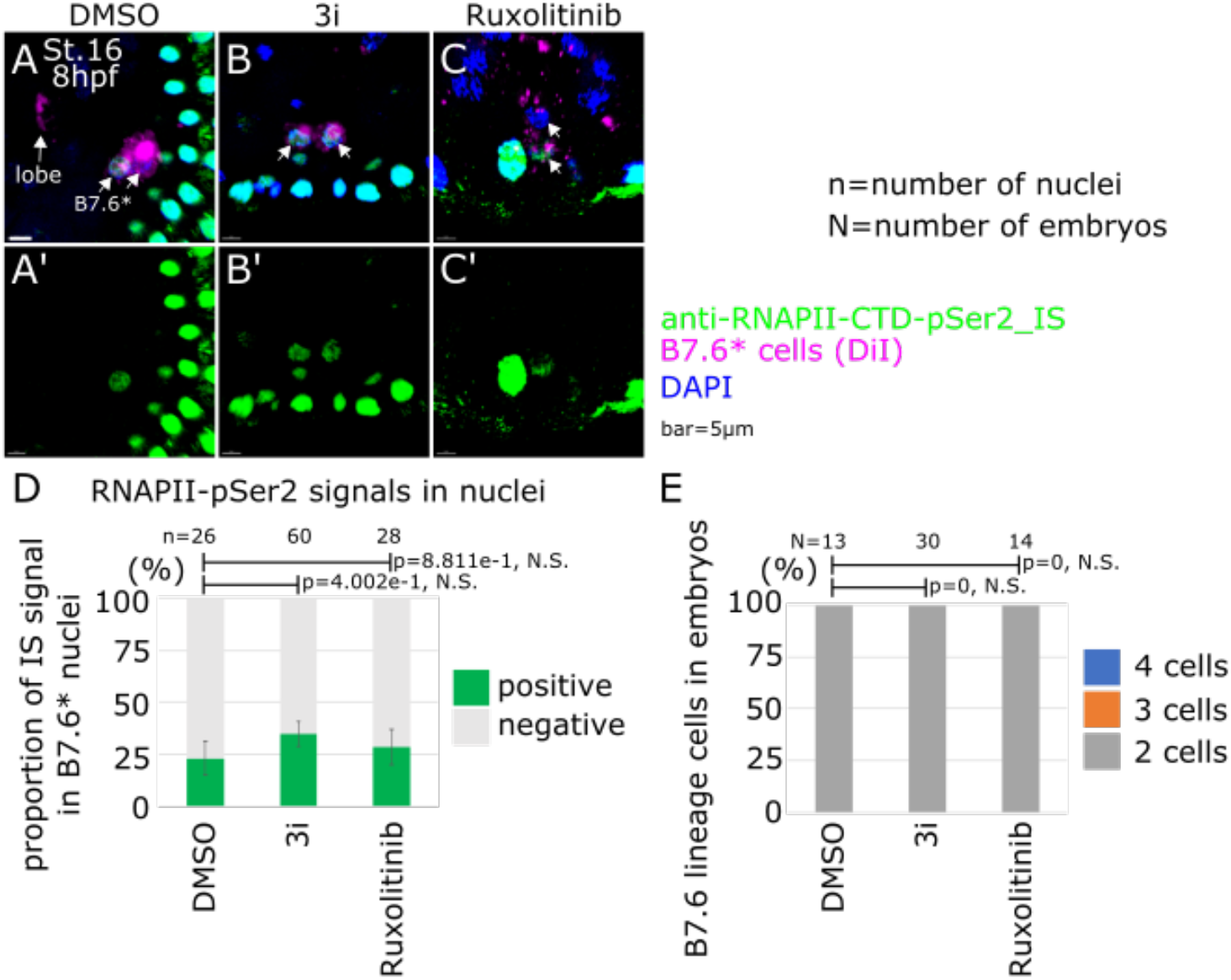
(A-C) Immunostaining was done with anti-RNAPII-CTD-pSer2 antibody under each pharmacological inhibitor treatment at 8hpf. Scale bar=5μm. (D) Proportion of signals of immunostaining with anti-RNAPII-CTD-pSer2 antibody in nuclei under each pharmacological inhibitor treatment. (E) Proportion of cell numbers of B7.6* cells in embryos under each pharmacological inhibitor treatment. Error bars indicate standard error. p-value was calculated by z-test. p>0.05; N.S, 0.05>p>0.01; *, 0.01>p; **.

## Footnotes

1 As we show in this study, B7.6 cells do not actually divide, and the previously named B8.11 cell is actually a cellular fragment, which we herein call the lobe, by analogy with a phenomenon described in *C. elegans*.

## References

Abdu Y, Maniscalco C, Heddleston JM, Chew T-L & Nance J (2016) Developmentally programmed germ cell remodelling by endodermal cell cannibalism. Nat Cell Biol 18: 1302–1310

Anderson R, Copeland TK, Schöler H, Heasman J & Wylie C (2000) The onset of germ cell migration in the mouse embryo. Mech Dev 91: 61–68

Batchelder C, Dunn MA, Choy B, Suh Y, Cassie C, Shim EY, Shin TH, Mello C, Seydoux G & Blackwell TK (1999) Transcriptional repression by the Caenorhabditis elegans germ-line protein PIE-1. Genes Dev 13: 202–212

Bensaude O (2011) Inhibiting eukaryotic transcription: Which compound to choose? How to evaluate its activity? Transcription 2: 103–108

Bhartiya D, Anand S, Patel H & Parte S (2017) Making gametes from alternate sources of stem cells: past, present and future. Reprod Biol Endocrinol 15: 89

Blackwell TK (2004) Germ cells: finding programs of mass repression. Curr Biol 14: R229–30

Brozovic M, Dantec C, Dardaillon J, Dauga D, Faure E, Gineste M, Louis A, Naville M, Nitta KR, Piette J, et al (2018) ANISEED 2017: extending the integrated ascidian database to the exploration and evolutionary comparison of genome-scale datasets. Nucleic Acids Res 46: D718–D725

Brunetti R, Gissi C, Pennati R, Caicci F, Gasparini F & Manni L (2015) Morphological evidence that the molecularly determined *Ciona intestinalis* type A and type B are different species: *Ciona robusta* and *Ciona intestinalis*. J Zoolog Syst Evol Res 53: 186–193

Chenevert J, Pruliere G, Ishii H, Sardet C & Nishikata T (2013) Purification of mitochondrial proteins HSP60 and ATP synthase from ascidian eggs: implications for antibody specificity. PLoS One 8: e52996

Christiaen L, Wagner E, Shi W & Levine M (2009a) Isolation of sea squirt (Ciona) gametes, fertilization, dechorionation, and development. Cold Spring Harb Protoc 2009: db.prot5344

Christiaen L, Wagner E, Shi W & Levine M (2009b) Whole-mount in situ hybridization on sea squirt (Ciona intestinalis) embryos. Cold Spring Harb Protoc 2009: db.prot5348

Dardaillon J, Dauga D, Simion P, Faure E, Onuma TA, DeBiasse MB, Louis A, Nitta KR, Naville M, Besnardeau L, et al (2020) ANISEED 2019: 4D exploration of genetic data for an extended range of tunicates. Nucleic Acids Res 48: D668–D675

Davidson EH & Levine MS (2008) Properties of developmental gene regulatory networks. Proc Natl Acad Sci U S A 105:20063–20066

Dehal P, Satou Y, Campbell RK, Chapman J, Degnan B, De Tomaso A, Davidson B, Di Gregorio A, Gelpke M, Goodstein DM, et al (2002) The draft genome of Ciona intestinalis: insights into chordate and vertebrate origins. Science 298: 2157–2167

D’Orazio FM, Balwierz PJ, González AJ, Guo Y, Hernández-Rodríguez B, Wheatley L, Jasiulewicz A, Hadzhiev Y, Vaquerizas JM, Cairns B, et al (2021) Germ cell differentiation requires Tdrd7-dependent chromatin and transcriptome reprogramming marked by germ plasm relocalization. Dev Cell 56: 641–656.e5

Dröscher A (2014) Images of cell trees, cell lines, and cell fates: the legacy of Ernst Haeckel and August Weismann in stem cell research. Hist Philos Life Sci 36: 157–186

Feinberg S, Roure A, Piron J & Darras S (2019) Antero-posterior ectoderm patterning by canonical Wnt signaling during ascidian development. PLoS Genet 15: e1008054

Gafni O, Weinberger L, Mansour AA, Manor YS, Chomsky E, Ben-Yosef D, Kalma Y, Viukov S, Maza I, Zviran A, et al (2013) Derivation of novel human ground state naive pluripotent stem cells. Nature 504: 282–286

Gline S, Kaplan N, Bernadskaya Y, Abdu Y & Christiaen L (2015) Surrounding tissues canalize motile cardiopharyngeal progenitors towards collective polarity and directed migration. Development 142: 544–554 doi:10.1242/dev.115444 [PREPRINT]

Graf T & Enver T (2009) Forcing cells to change lineages. Nature 462: 587–594

Gregor T, Garcia HG & Little SC (2014) The embryo as a laboratory: quantifying transcription in Drosophila. Trends Genet 30: 364–375

Hanna J, Cheng AW, Saha K, Kim J, Lengner CJ, Soldner F, Cassady JP, Muffat J, Carey BW & Jaenisch R (2010) Human embryonic stem cells with biological and epigenetic characteristics similar to those of mouse ESCs. Proc Natl Acad Sci U S A 107: 9222–9227

Hanyu-Nakamura K, Sonobe-Nojima H, Tanigawa A, Lasko P & Nakamura A (2008) Drosophila Pgc protein inhibits P-TEFb recruitment to chromatin in primordial germ cells. Nature 451: 730–733

Harder M, Reeves W, Byers C, Santiago M & Veeman M (2019) Multiple inputs into a posterior-specific regulatory network in the Ciona notochord. Dev Biol 448: 136–146

Hayashi K & Saitou M (2014) Perspectives of germ cell development in vitro in mammals. Animal Science Journal 85: 617–626 doi:10.1111/asj.12199 [PREPRINT]

Hino K, Satou Y, Yagi K & Satoh N (2003) A genomewide survey of developmentally relevant genes in Ciona intestinalis. VI. Genes for Wnt, TGFbeta, Hedgehog and JAK/STAT signaling pathways. Dev Genes Evol 213: 264–272

Hotta K, Mitsuhara K, Takahashi H, Inaba K, Oka K, Gojobori T & Ikeo K (2007) A web-based interactive developmental table for the ascidian Ciona intestinalis, including 3D real-image embryo reconstructions: I. From fertilized egg to hatching larva. Dev Dyn 236:1790–1805

Hu M, Krause D, Greaves M, Sharkis S, Dexter M, Heyworth C & Enver T (1997) Multilineage gene expression precedes commitment in the hemopoietic system. Genes Dev 11: 774–785

Hudson C, Darras S, Caillol D, Yasuo H & Lemaire P (2003) A conserved role for the MEK signalling pathway in neural tissue specification and posteriorisation in the invertebrate chordate, the ascidian Ciona intestinalis. Development 130: 147–159

Imai KS (2004) Gene expression profiles of transcription factors and signaling molecules in the ascidian embryo: towards a comprehensive understanding of gene networks. Development 131: 4047–4058 doi:10.1242/dev.01270 [PREPRINT]

Imai KS, Levine M, Satoh N & Satou Y (2006) Regulatory blueprint for a chordate embryo. Science 312: 1183–1187

Jostes S & Schorle H (2018) Signals and transcription factors for specification of human germ cells. Stem Cell Investig 5: 13

Kanamori M, Oikawa K, Tanemura K & Hara K (2019) Mammalian germ cell migration during development, growth, and homeostasis. Reprod Med Biol 18: 247–255

Kang MK & Han SJ (2011) Post-transcriptional and post-translational regulation during mouse oocyte maturation. BMB Rep 44: 147–157

Karaiskou A, Swalla BJ, Sasakura Y & Chambon J-P (2015) Metamorphosis in solitary ascidians. Genesis 53: 34–47

Kawai N, Ogura Y, Ikuta T, Saiga H, Hamada M, Sakuma T, Yamamoto T, Satoh N & Sasakura Y (2015) Hox10-regulated endodermal cell migration is essential for development of the ascidian intestine. Dev Biol 403: 43–56

Krasovec G, Robine K, Quéinnec E, Karaiskou A & Chambon JP (2019) Ci-hox12 tail gradient precedes and participates in the control of the apoptotic-dependent tail regression during Ciona larva metamorphosis. Dev Biol 448: 237–246

Kumano G, Takatori N, Negishi T, Takada T & Nishida H (2011) A maternal factor unique to ascidians silences the germline via binding to P-TEFb and RNAP II regulation. Curr Biol 21: 1308–1313

Kutschera U & Niklas KJ (2004) The modern theory of biological evolution: an expanded synthesis. Naturwissenschaften 91: 255–276

Lawson KA, Dunn NR, Roelen BA, Zeinstra LM, Davis AM, Wright CV, Korving JP & Hogan BL (1999) Bmp4 is required for the generation of primordial germ cells in the mouse embryo. Genes Dev 13: 424–436

Lebedeva LA, Yakovlev KV, Kozlov EN, Schedl P, Deshpande G & Shidlovskii YV (2018) Transcriptional quiescence in primordial germ cells. Crit Rev Biochem Mol Biol 53: 579–595

Lehmann R (2012) Germline Stem Cells: Origin and Destiny. Cell Stem Cell 10: 729–739 doi:10.1016/j.stem.2012.05.016 [PREPRINT]

Levine M & Davidson EH (2005) Gene regulatory networks for development. Proc Natl Acad Sci U S A 102: 4936–4942

Levine M & Tjian R (2003) Transcription regulation and animal diversity. Nature 424: 147–151

Little SC, Sinsimer KS, Lee JJ, Wieschaus EF & Gavis ER (2015) Independent and coordinate trafficking of single Drosophila germ plasm mRNAs. Nat Cell Biol 17: 558–568

Mainpal R, Nance J & Yanowitz JL (2015) A germ cell determinant reveals parallel pathways for germ line development in Caenorhabditis elegans. Development 142: 3571–3582

Malfant M, Coudret J, Le Merdy R & Viard F (2017) Effects of temperature and salinity on juveniles of two ascidians, one native and one invasive, and their hybrids. J Exp Mar Bio Ecol 497: 180–187

Maniscalco C, Hall AE & Nance J (2020) An interphase contractile ring reshapes primordial germ cells to allow bulk cytoplasmic remodeling. J Cell Biol 219

McIntyre DC & Nance J (2020) Niche Cell Wrapping Ensures Primordial Germ Cell Quiescence and Protection from Intercellular Cannibalism. Curr Biol 30: 708–714.e4

Mello CC, Schubert C, Draper B, Zhang W, Lobel R & Priess JR (1996) The PIE-1 protein and germline specification in C. elegans embryos. Nature 382: 710–712 doi:10.1038/382710a0 [PREPRINT]

Mitsunaga S & Shioda T (2018) Evolutionarily diverse mechanisms of germline specification among mammals: what about us? Stem Cell Investig 5: 12

Miyaoku K, Nakamoto A, Nishida H & Kumano G (2018) Control of Pem protein level by localized maternal factors for transcriptional regulation in the germline of the ascidian, Halocynthia roretzi. PLoS One 13: e0196500

Moris N, Pina C & Arias AM (2016) Transition states and cell fate decisions in epigenetic landscapes. Nat Rev Genet 17:693–703

Nakamura A, Amikura R, Mukai M, Kobayashi S & Lasko PF (1996) Requirement for a Noncoding RNA in Drosophila Polar Granules for Germ Cell Establishment. Science 274: 2075–2079 doi:10.1126/science.274.5295.2075 [PREPRINT]

Nakamura A & Seydoux G (2008) Less is more: specification of the germline by transcriptional repression. Development 135: 3817–3827

Nimmo RA, May GE & Enver T (2015) Primed and ready: understanding lineage commitment through single cell analysis. Trends Cell Biol 25: 459–467

Nishida H & Satoh N (1983) Cell lineage analysis in ascidian embryos by intracellular injection of a tracer enzyme. I. Up to the eight-cell stage. Dev Biol 99: 382–394

Nishikata T, Yamada L, Mochizuki Y, Satou Y, Shin-i T, Kohara Y & Satoh N (2001) Profiles of maternally expressed genes in fertilized eggs of Ciona intestinalis. Dev Biol 238: 315–331

Oda-Ishii I, Kubo A, Kari W, Suzuki N, Rothbächer U & Satou Y (2016) A Maternal System Initiating the Zygotic Developmental Program through Combinatorial Repression in the Ascidian Embryo. PLoS Genet 12: e1006045

Ohinata Y, Ohta H, Shigeta M, Yamanaka K, Wakayama T & Saitou M (2009) A signaling principle for the specification of the germ cell lineage in mice. Cell 137: 571–584

Ohta N, Kaplan N, Ng JT, Gravez BJ & Christiaen L (2020) Asymmetric Fitness of Second-Generation Interspecific Hybrids Between Ciona robusta and Ciona intestinalis. G3 (Bethesda) 10: 2697–2711

Ohta N & Satou Y (2013) Multiple signaling pathways coordinate to induce a threshold response in a chordate embryo. PLoS Genet 9: e1003818

Olsen LC, Kourtesis I, Busengdal H, Jensen MF, Hausen H & Chourrout D (2018) Evidence for a centrosome-attracting body like structure in germ-soma segregation during early development, in the urochordate Oikopleura dioica. BMC Dev Biol 18: 4

Paix A, Yamada L, Dru P, Lecordier H, Pruliere G, Chenevert J, Satoh N & Sardet C (2009) Cortical anchorages and cell type segregations of maternal postplasmic/PEM RNAs in ascidians. Dev Biol 336: 96–111

Pilato G, Pilato G, D’Urso V, Viglianisi F, Sammartano F, Sabella G & Lisi O (2013) The problem of the origin of primordial germ cells (PGCs) in vertebrates: historical review and a possible solution. The International Journal of Developmental Biology 57: 809–819 doi:10.1387/ijdb.120261gp [PREPRINT]

Prodon F, Yamada L, Shirae-Kurabayashi M, Nakamura Y & Sasakura Y (2007) Postplasmic/PEM RNAs: a class of localized maternal mRNAs with multiple roles in cell polarity and development in ascidian embryos. Dev Dyn 236: 1698–1715

Razy-Krajka F, Gravez B, Kaplan N, Racioppi C, Wang W & Christiaen L (2018) An FGF-driven feed-forward circuit patterns the cardiopharyngeal mesoderm in space and time. Elife 7

Razy-Krajka F, Lam K, Wang W, Stolfi A, Joly M, Bonneau R & Christiaen L (2014) Collier/OLF/EBF-dependent transcriptional dynamics control pharyngeal muscle specification from primed cardiopharyngeal progenitors. Dev Cell 29: 263–276

Ristoratore F, Spagnuolo A, Aniello F, Branno M, Fabbrini F & Di Lauro R (1999) Expression and functional analysis of Cititf1, an ascidian NK-2 class gene, suggest its role in endoderm development. Development 126: 5149–5159

Robert VJ, Garvis S & Palladino F (2015) Repression of somatic cell fate in the germline. Cell Mol Life Sci 72: 3599–3620

Sardet C, Paix A, Prodon F, Dru P & Chenevert J (2007) From oocyte to 16-cell stage: cytoplasmic and cortical reorganizations that pattern the ascidian embryo. Dev Dyn 236: 1716–1731

Sasakura Y, Ogasawara M & Makabe KW (2000) Two pathways of maternal RNA localization at the posterior-vegetal cytoplasm in early ascidian embryos. Dev Biol 220: 365–378

Sato A, Shimeld SM & Bishop JDD (2014) Symmetrical reproductive compatibility of two species in the Ciona intestinalis (Ascidiacea) species complex, a model for marine genomics and developmental biology. Zoolog Sci 31: 369–374

Satou Y & Imai KS (2015) Gene regulatory systems that control gene expression in the Ciona embryo. Proc Jpn Acad Ser B Phys Biol Sci 91: 33–51

Satou Y, Imai KS & Satoh N (2004) The ascidian Mesp gene specifies heart precursor cells. Development 131: 2533–2541

Satou Y, Kawashima T, Shoguchi E, Nakayama A & Satoh N (2005) An integrated database of the ascidian, Ciona intestinalis: towards functional genomics. Zoolog Sci 22: 837–843

Schwartz AZA, Tsyba N, Abdu Y, Patel MR & Nance J (2022) Independent regulation of mitochondrial DNA quantity and quality in Caenorhabditis elegans primordial germ cells. Elife 11

Seydoux G & Braun RE (2006) Pathway to totipotency: lessons from germ cells. Cell 127: 891–904

Seydoux G & Dunn MA (1997) Transcriptionally repressed germ cells lack a subpopulation of phosphorylated RNA polymerase II in early embryos of Caenorhabditis elegans and Drosophila melanogaster. Development 124: 2191–2201

Shirae-Kurabayashi M, Matsuda K & Nakamura A (2011) Ci-Pem-1 localizes to the nucleus and represses somatic gene transcription in the germline of Ciona intestinalis embryos. Development 138: 2871–2881

Shirae-Kurabayashi M, Nishikata T, Takamura K, Tanaka KJ, Nakamoto C & Nakamura A (2006) Dynamic redistribution of vasa homolog and exclusion of somatic cell determinants during germ cell specification in Ciona intestinalis. Development 133: 2683–2693

Sobell HM (1985) Actinomycin and DNA transcription. Proc Natl Acad Sci U S A 82: 5328–5331

Strome S & Updike D (2015) Specifying and protecting germ cell fate. Nat Rev Mol Cell Biol 16: 406–416

Suzuki MM, Nishikawa T & Bird A (2005) Genomic approaches reveal unexpected genetic divergence within Ciona intestinalis. J Mol Evol 61: 627–635

Takamura K, Fujimura M & Yamaguchi Y (2002) Primordial germ cells originate from the endodermal strand cells in the ascidian Ciona intestinalis. Dev Genes Evol 212: 11–18

Takashima Y, Guo G, Loos R, Nichols J, Ficz G, Krueger F, Oxley D, Santos F, Clarke J, Mansfield W, et al (2015) Resetting Transcription Factor Control Circuitry toward Ground-State Pluripotency in Human. Cell 162: 452–453

Tang WWC, Kobayashi T, Irie N, Dietmann S & Surani MA (2016) Specification and epigenetic programming of the human germ line. Nat Rev Genet 17: 585–600

Tassy O, Dauga D, Daian F, Sobral D, Robin F, Khoueiry P, Salgado D, Fox V, Caillol D, Schiappa R, et al (2010) The ANISEED database: digital representation, formalization, and elucidation of a chordate developmental program. Genome Res 20:1459–1468

Tkačik G & Gregor T (2021) The many bits of positional information. Development 148

Tokuoka M, Kobayashi K, Lemaire P & Satou Y (2022) Protein kinases and protein phosphatases encoded in the Ciona robusta genome. Genesis 60: e23471

Trcek T & Lehmann R (2019) Germ granules in Drosophila. Traffic 20: 650–660

Waki K, Imai KS & Satou Y (2015) Genetic pathways for differentiation of the peripheral nervous system in ascidians. Nature Communications 6 doi:10.1038/ncomms9719 [PREPRINT]

Wang W, Niu X, Stuart T, Jullian E, Mauck WM 3rd, Kelly RG, Satija R & Christiaen L (2019) A single-cell transcriptional roadmap for cardiopharyngeal fate diversification. Nat Cell Biol 21: 674–686

Wang W, Razy-Krajka F, Siu E, Ketcham A & Christiaen L (2013) NK4 antagonizes Tbx1/10 to promote cardiac versus pharyngeal muscle fate in the ascidian second heart field. PLoS Biol 11: e1001725

Ware CB, Nelson AM, Mecham B, Hesson J, Zhou W, Jonlin EC, Jimenez-Caliani AJ, Deng X, Cavanaugh C, Cook S, et al (2014) Derivation of naive human embryonic stem cells. Proc Natl Acad Sci U S A 111: 4484–4489

Weismann A (1892) Das Keimplasma: eine Theorie der Vererbung

Wong T-T & Zohar Y (2015) Production of reproductively sterile fish by a non-transgenic gene silencing technology. Sci Rep 5: 15822

Yamada L (2006) Embryonic expression profiles and conserved localization mechanisms of pem/postplasmic mRNAs of two species of ascidian, Ciona intestinalis and Ciona savignyi. Dev Biol 296: 524–536

Yamada L, Kobayashi K, Satou Y & Satoh N (2005) Microarray analysis of localization of maternal transcripts in eggs and early embryos of the ascidian, Ciona intestinalis. Dev Biol 284: 536–550

Ying Q-L, Wray J, Nichols J, Batlle-Morera L, Doble B, Woodgett J, Cohen P & Smith A (2008) The ground state of embryonic stem cell self-renewal. Nature 453: 519–523

Ying, Ying Y, Qi X & Zhao G-Q (2002) Induction of Primordial Germ Cells from Pluripotent Epiblast. The Scientific World JOURNAL 2: 801–810 doi:10.1100/tsw.2002.155 [PREPRINT]

Yoshida S, Marikawa Y & Satoh N (1996) Posterior end mark, a novel maternal gene encoding a localized factor in the ascidian embryo. Development 122: 2005–2012 doi:10.1242/dev.122.7.2005 [PREPRINT]

Yu L, Wei Y, Sun H-X, Mahdi AK, Pinzon Arteaga CA, Sakurai M, Schmitz DA, Zheng C, Ballard ED, Li J, et al (2020) Derivation of Intermediate Pluripotent Stem Cells Amenable to Primordial Germ Cell Specification. Cell Stem Cell doi:10.1016/j.stem.2020.11.003 [PREPRINT]

