## Supplementary material for "Cellular remodeling and JAK inhibition promote zygotic gene expression in the *Ciona* germline": Table S1 (from another version of this preprint: version 2, 2021-11-09)

### Supplemental Table.S1 Primers

| Gene name | Type | | Primers sequence |
|---|---|---|---|
| *Mef2* | intron | Forward | 5'-ACTCCATGCCCACTTACGAA-3' |
| | | Reverse | 5'-GGATCGCGATGTAAAGGACG-3' |
| | intron | Forward | 5'-CGACACACCCGTTAATCTGG-3' |
| | | Reverse | 5'-GGGGATGTAGTCTGAAGGGG-3' |
| *MS4A15/2* | exon | Forward | 5'-CAAAGGAAATGTCAGTTAC-3' |
| | | Reverse | 5'-GTTCCAGAAACTAAATAAG-3' |
| *Pem-1* | exon | Forward | 5'-AAAGAAAGTGCATAAATATG-3' |
| | | Reverse | 5'-AATCGTGAAAAAGTACTAAC-3' |
